# Bacterial antiviral defense pathways encode eukaryotic-like ubiquitination systems

**DOI:** 10.1101/2023.09.26.559546

**Authors:** Lydia R. Chambers, Qiaozhen Ye, Jiaxi Cai, Minheng Gong, Hannah E. Ledvina, Huilin Zhou, Aaron T. Whiteley, Raymond T. Suhandynata, Kevin D. Corbett

## Abstract

Ubiquitination and related pathways play crucial roles in protein homeostasis, signaling, and innate immunity^1-3^. In these pathways, an enzymatic cascade of E1, E2, and E3 proteins conjugates ubiquitin or a ubiquitin-like protein (Ubl) to targetprotein lysine residues^4^. Bacteria encode ancient relatives of E1 and Ubl proteins involved in sulfur metabolism^5,6^ but these proteins do not mediate Ubl-target conjugation, leaving open the question of whether bacteria can perform ubiquitination-like protein conjugation. Here, we demonstrate that a bacterial antiviral immune system encodes a complete ubiquitination pathway. Two structures of a bacterial E1:E2:Ubl complex reveal striking architectural parallels with canonical eukaryotic ubiquitination machinery. The bacterial E1 encodes an N-terminal inactive adenylation domain (IAD) and a C-terminal active adenylation domain (AAD) with a mobile α-helical insertion containing the catalytic cysteine (CYS domain). One structure reveals a pre-reaction state with the bacterial Ubl C-terminus positioned for adenylation, and the E1 CYS domain poised nearby for thioester formation. A second structure mimics an E1-to-E2 transthioesterification state, with the E1 CYS domain rotated outward and its catalytic cysteine adjacent to the bound E2. We show that a deubiquitinase (DUB) in the same pathway pre-processes the bacterial Ubl, exposing its C-terminal glycine for adenylation. Finally, we show that the bacterial E1 and E2 collaborate to conjugate Ubl to target-protein lysine residues. Together, these data reveal that bacteria possess *bona fide* ubiquitination systems with strong mechanistic and architectural parallels to canonical eukaryotic ubiquitination pathways, suggesting that these pathways arose first in bacteria.

## Introduction

The conjugation of ubiquitin and Ubls to target proteins involves three enzymes termed E1, E2, and E3. An E1 “activating protein” catalyzes the formation of a high-energy acyl adenylate intermediate through the reaction of ATP with the Ubl C-terminus, then a cysteine on the E1 attacks that intermediate to form a thioester link to the Ubl^4^. Next, a cysteine on an E2 “carrier protein” attacks the E1∼Ubl thioester to form a second E2∼Ubl thioester. Finally, conjugation of the Ubl to a lysine side chain on a target protein is mediated by an E3 “ligase,” which may simply act as an E2-target adapter (as in RING-family E3s) or form a third cysteine∼Ubl thioester intermediate before transfer to a target (as in HECT-family E3s). In all Ubl conjugation pathways, specific peptidases termed deubiquitinases or DUBs cleave Ubl-lysine isopeptide linkages in addition to pre-processing many Ubls to expose their reactive C-terminal glycine residue for E1-mediated catalysis^7^.

Bacteria encode widespread E1-like adenylation enzymes (ThiF, MoeB) that act on ubiquitin-like proteins (ThiS, MoaD), but these pathways do not encode E2-like proteins and do not mediate Ubl-target conjugation^5,6^. Over the past two decades, sparsely distributed bacterial operons have been reported that encode distinct combinations of putative E1, E2, Ubl, and DUB proteins^8-11^, plus rare examples of RING-type E3 proteins^12^. One such operon family, termed Bil (<u>B</u>acterial ISG15-like gene) encodes putative E1, E2, Ubl, and DUB proteins and protects its host against bacteriophage (phage) infection by specifically modifying a virion structural protein^13,14^. These findings suggest that, counter to prevailing models in which complete ubiquitination pathways arose first in archaea^15,16^, bacteria encode ancient ubiquitination-like pathways that participate in antiviral defense. To date, however, Ubl-target conjugation has not been demonstrated in these pathways.

## Results

### Identification of bacterial operons encoding a complete ubiquitination pathway

We recently showed that bacterial Type II CBASS immune systems encode an E1-E2 fusion protein (Cap2) that resembles the noncanonical eukaryotic E1 Atg7 and its partner E2s Atg3 and Atg10 ^17^. Cap2 conjugates a cGAS (cyclic GMP-AMP synthase)-like protein to target proteins using a ubiquitination-like mechanism^17,18^. We additionally identified four bacterial operon families that encode different combinations of E1, E2, JAB-family DUB, and Ubl proteins, one of which encoded a multidomain E1 protein and an uncharacterized protein termed CEHH after four conserved residues (cysteine, glutamate, histidine, histidine)^17^. In an earlier bioinformatic analysis of prokaryotic ubiquitination-related proteins, this operon family was denoted “6C” and described as encoding three proteins: multidomain E1, CEHH, and a JAB-family DUB^8^. Our sequence analysis revealed that CEHH likely represents a diverged E2-like protein, and further that some CEHH-containing operons also encode a Ubl. Together, these observations suggested that the CEHH/6C operons may represent a complete ancestral ubiquitination system.

We performed comprehensive sequence searches using a multidomain E1 protein from *Ensifer aridi* TW10 and identified 162 nonredundant CEHH/6C operons, with most instances in the plant-associated Rhizobiaceae family (Figure 1a, Supplementary Table 1). Most of these operons (135/162, 83%) encode four proteins: E1, CEHH/E2, DUB, and Ubl. We observed that many CEHH/6C operons are located near CRISPR/Cas and other known immune systems, suggesting that they occur in “defense islands” enriched for antiviral and other conflict systems^19-22^. Twenty-five CEHH/6C operons (15%) are associated with CapH/CapP transcriptional regulators, which we previously found to be associated with diverse bacterial immune systems and which activate these systems’ expression in response to DNA damage^23^. Together, these data show that CEHH/6C operons encode a complete ubiquitination-like pathway with predicted E1, E2, Ubl, and DUB proteins, and further suggest that these operons may participate in antiviral defense.

**Figure 1.**
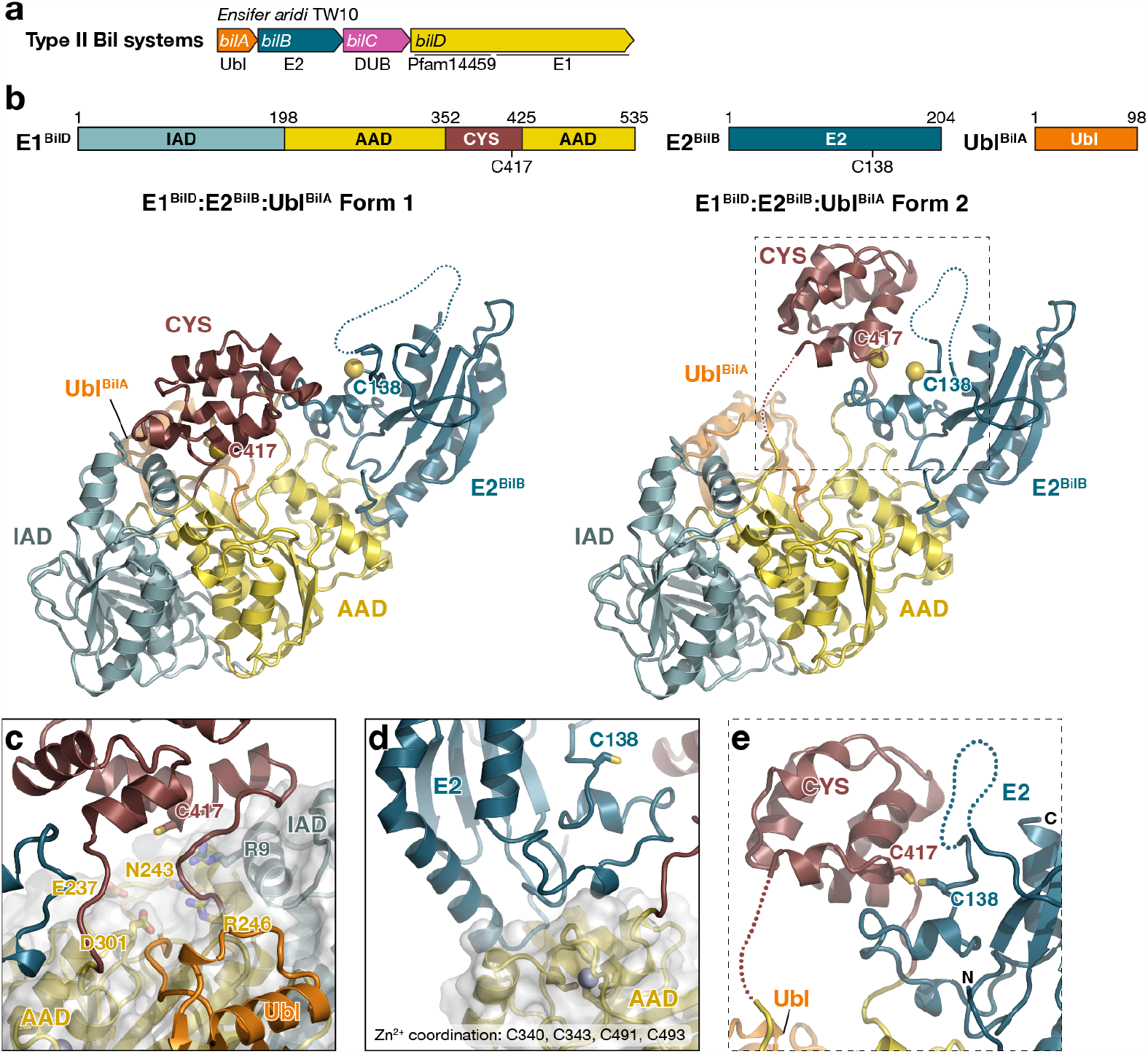
Structure of a bacterial E1-E2-Ubl complex. **(a)** Schematic of a Type II BilABCD operon from *Ensifer aridi* TW10. See **Supplementary Table 1** for a full list of Type II BilABCD operons, and **Supplementary Figure 1** for comparison with Type I BilABCD operons. **(b)** *Top:* Domain schematic of *E. aridi* E1^BilD^, E2^BilB^, and Ubl^BilA^. The inactive adenylation domain (IAD) of E1^BilD^ is colored light blue, the active adenylation domain (AAD) yellow, and the α-helical insertion containing the putative catalytic cysteine (CYS domain) brown. Putative catalytic cysteine residues in E1^BilD^ (C417) and E2^BilB^ (C138) are noted. *Left: E. aridi* E1^BilD^:E2^BilB^:Ubl^BilA^ complex Form 1 crystal structure, with domains colored as in the schematic. Putative catalytic cysteine residues are shown as yellow spheres. *Right: E. aridi* E1^BilD^:E2^BilB^:Ubl^BilA^ complex Form 2 crystal structure. See **Supplementary Figure 3** for comparison to a canonical eukaryotic E1(NAE1-UBA3):E2(Ubc12):Ubl(NEDD8) complex. **(c)** Closeup view of the Ubl^BilA^ C-terminus (orange) docked in the E1BilD adenylation active site (yellow, with conserved active-site residues shown as sticks and labeled). The E1 catalytic cysteine residue (C417) is positioned 14 Å away from the Ubl C-terminus. See **Supplementary Figure 3c, g** for comparison of bacterial and eukaryotic E1 adenylation active sites. **(d)** Closeup view of the E1^BilD^-E2^BilB^ binding interface, including the structural Zn^2+^ ion coordinated by four cysteine residues in E1^BilD^. See **Supplementary Figure 3d, h** for comparison of bacterial and eukaryotic E1-E1 interfaces. **(e)** Closeup view of the E1^BIlD^ and E2^BilB^ catalytic cysteine positions in the Form 2 structure. Compared to Form 1, the E1 CYS domain is rotated upward and the E1^BilD^ catalytic cysteine (C417) is positioned within 2 Å of the E2 catalytic cysteine (C138), mimicking the structural state adopted during an E1^BilD^-to-E2^BilB^ transthioesterification reaction.

We observed that the gene order in CEHH/6C operons (Ubl–CEHH/E2–DUB–E1) matches that of BilABCD operons (Ubl–E2–DUB–E1; Supplementary Figure 1a)^13^. Sequence alignments of the four equivalent protein groups in CEHH/6C and BilABCD operons show that in each case, these proteins share identifiable sequence homology but segregate into distinct groups in unrooted evolutionary trees (Supplementary Figure 1b-d). The broad parallels between CEHH and BilABCD operons, combined with these specific differences, prompted us to rename the CEHH/6C operons as Type II BilABCD operons.

### Structure of a bacterial E1-E2-Ubl complex

We cloned and expressed the four proteins from a Type II BilABCD operon from *Ensifer aridi* TW10 (Ubl^BilA^, E2^BilB^, DUB^BilC^, and E1^BilD^) for biochemical and structural analysis. While both E1^BilD^ and E2^BilB^ were insoluble when overexpressed in *E. coli*, they formed a soluble complex when coexpressed with Ubl^BilA^. The three-protein E1^BilD^:E2^BilB^:Ubl^BilA^ complex eluted from a size exclusion chromatography column at a position consistent with a ∼250 kDa complex (Stokes radius 57.6 Å), suggesting that it forms a 2:2:2 complex (180.4 kDa) or a higher-order oligomer (Supplementary Figure 2). We crystallized and determined two independent X-ray crystal structures of the E1^BilD^:E2^BilB^:Ubl^BilA^ complex to 2.5 Å resolution (Form 1) and 2.7 Å resolution (Form 2) (Figure 1b, Supplementary Table 2).

E1^BilD^ encodes an N-terminal domain of unknown function (Pfam14459) annotated as “prokaryotic E2 family C” and a C-terminal E1-like adenylation domain (Supplementary Figure 1a), and was originally speculated to comprise an E2-E1 fusion protein^8^. Our structure shows that E1^BilD^ adopts an overall architecture strikingly similar to that of canonical eukaryotic E1 proteins (Figure 1b, Figure 2a-b, Supplementary Figure 3). Whereas all known bacterial E1 and E1-like proteins including ThiS, MoeB, and Cap2 form homodimers through their adenylation domains^5,6,17^, E1^BilD^ possesses an N-terminal inactive adenylation domain (IAD) and a C-terminal active adenylation domain (AAD) that form a structurally related pseudo-dimer. Inserted into the “crossover loop” of the E1^BilD^ AAD is an ∼80 amino acid α-helical domain containing a conserved cysteine residue (CYS domain). This domain architecture – IAD, AAD, and CYS – is a hallmark of canonical eukaryotic E1 proteins. The closest structural relatives of E1^BilD^ are the human ubiquitin/FAT10 E1 UBA6 (Cα r.m.s.d. 3.5 Å over 401 residue pairs spanning IAD and AAD)^24^ and the heterodimeric human NEDD8 E1 NAE1-UBA3 (Cα r.m.s.d. 3.6 Å over 221 residue pairs between E1^BilD^ IAD and NAE1 IAD, and Cα r.m.s.d. 3.7 Å over 250 residue pairs between E1^BilD^ AAD and UBA3 AAD)^25^ (Figure 2a-b, Supplementary Figure 3). Like these eukaryotic E1s, E1^BilD^ possesses a single active adenylation site in the AAD, with one key arginine residue (Arg9) provided by the IAD (Figure 1c, Supplementary Figure 3c, g). The E1^BilD^ CYS domain is linked to the AAD via flexible linkers and is highly mobile, showing high B-factors compared to the IAD/AAD and adopting different positions in Form 1 versus Form 2 structures. In the Form 1 structure, the E1 conserved cysteine residue (Cys417) is poised directly above the adenylation active site, whereas in Form 2 this domain is rotated upward to contact the bound E2^BilB^ (see below). E1^BilD^ lacks other major structural features of canonical ubiquitin E1 proteins, including the first catalytic cysteine half-domain (FCCH) or coiled-coil domain (CC) that is often inserted within the IAD, and the C-terminal ubiquitin fold domain (UFD) that mediates E2 binding. E1^BilD^ coordinates a zinc ion through four cysteine residues, two in the AAD crossover loop and two near the protein’s C-terminus (Figure 1d); similar structural zinc ions are observed in SUMO and NEDD8 E1 proteins (Supplementary Figure 3d,h)^4,25,26^. Overall, the structural features of E1^BilD^ show surprising parallels with canonical eukaryotic E1 proteins that suggest a common ancestry.

**Figure 2.**
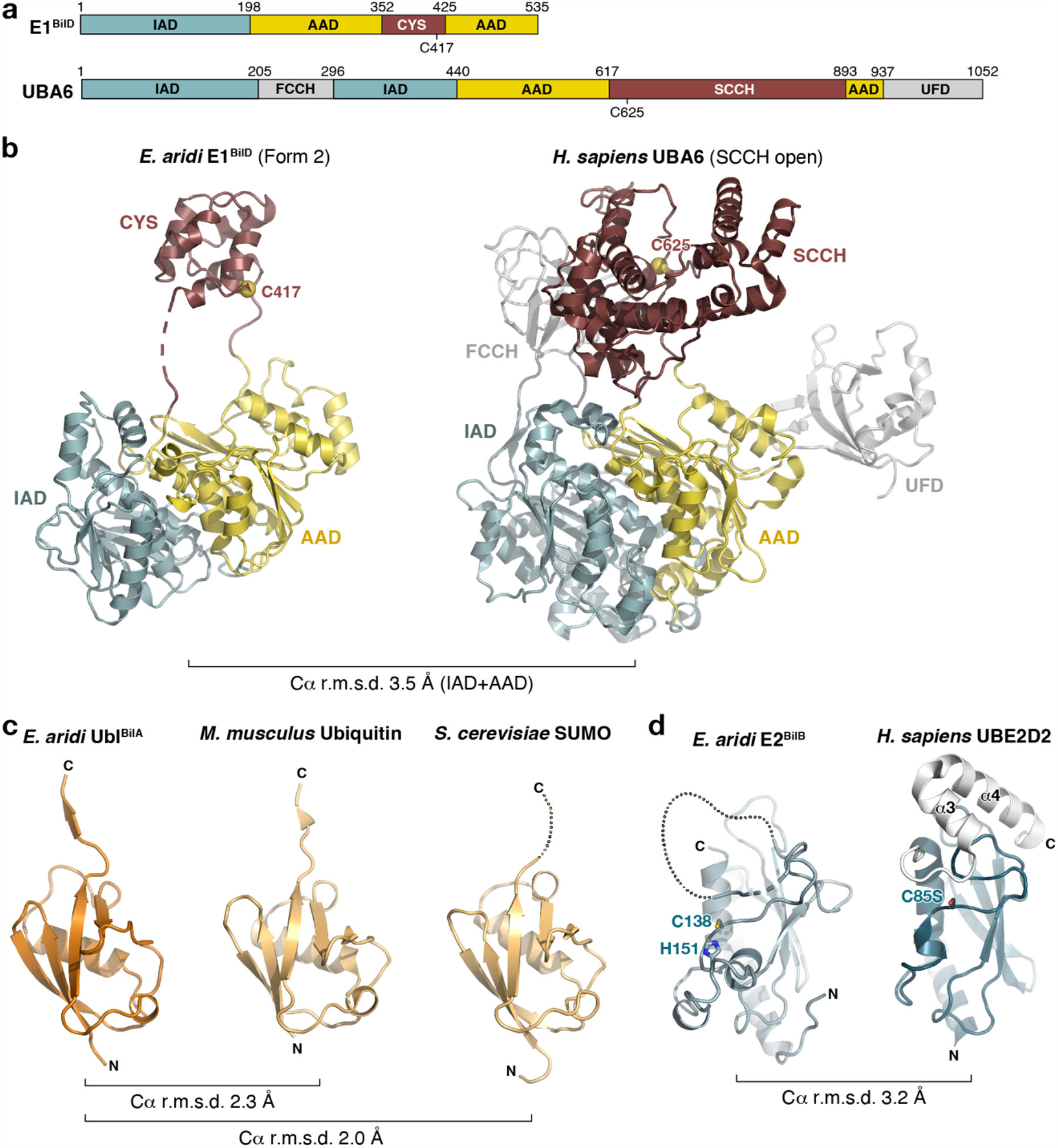
*E. aridi* E1^BilD^, E2^BilB^, and Ubl^BilA^ resemble canonical eukaryotic ubiquitination machinery. **(a)** Domain schematic of *E. aridi* E1^BilD^ and the *H. sapiens* ubiquitin/FAT10 E1 UBA6 (PDB ID 7SOL)^24^. IAD: inactive adenylation domain; AAD: active adenylation domain; CYS: catalytic cysteine-containing domain; FCCH: first catalytic cysteine half-domain; SCCH: second catalytic cysteine half-domain (equivalent to CYS); UFD: ubiquitin fold domain. **(b)** Structures of *E. aridi* E1^BilD^ (left) and *H. sapiens* UBA6 (PDB ID 7SOL)^24^, with domains colored as in panel (a) and catalytic cysteines shown as spheres and colored yellow. The Cα r.m.s.d. is 3.5 Å over 401 residue pairs spanning the two proteins’ IAD and AAD domains. **(c)** Structures of *E. aridi* Ubl^BilA^ (left), *M. musculus* ubiquitin (center; PDB ID 4NQL)^53^, and *S. cerevisiae* SUMO/Smt3 (right; PDB ID 5JNE)^28^. **(d)** Structures of *E. aridi* E2^BilB^ (left) and the *H. sapiens* E2 UBE2D2 (right; PDB ID 4DGG)^32^. Each protein’s catalytic cysteine residue is shown as sticks and labeled (Cys85 is mutated to serine in the *H. sapiens* UBE2D2 structure. *E. aridi* E2^BilB^ His151 is also shown as sticks. See **Supplementary Figure 5a** for an equivalent view showing all four conserved residues of the originally identified CEHH motif. For UBE2D2, the C-terminal α-helices not shared by E2^BilB^ are shown in white.

In contrast to identified Type I BilABCD operons whose Ubl (BilA) contains two predicted ubiquitin-like β-grasp domains^13^, *E. aridi* Ubl^BilA^ contains a single β-grasp domain (Figure 2c). The closest structural homologs to Ubl^BilA^ are mammalian ubiquitin (Cα r.m.s.d. 1.7 Å over 72 residue pairs)^27^, *S. cerevisiae* SUMO (Smt3; Cα r.m.s.d. 1.7 Å over 73 residue pairs)^28^, an archaeal ubiquitin-like protein (*Caldiarchaeum subterraneum* Ub; Cα r.m.s.d. 2.2 Å over 73 residue pairs)^29^, and ubiquitin-like domains in *H. sapiens* Talin2 (Cα r.m.s.d. 2.3 Å over 82 residue pairs)^30^ and *S. pombe* Rad60 (Cα r.m.s.d. 1.6 Å over 72 residue pairs)^31^. Ubl^BilA^ shows markedly less structural similarity to bacterial β-grasp proteins involved in sulfur metabolism, including ThiS (Cα r.m.s.d. 3.3 Å over 57 residue pairs)^6^ and MoaD (Cα r.m.s.d. 3.3 Å over 64 residue pairs)^5^. While *E. aridi* Ubl^BilA^ encodes a single β-grasp domain, Ubl^BilA^ proteins across Type II BilABCD operons show remarkable structural diversity: we identified examples of Ubls encoding up to three tandem β-grasp domains, and determined a crystal structure of one three-domain Ubl^BilA^ from *Methylobacterium brachiatum* DSM 19569 (Supplementary Figure 4a-d). We also identified Ubls with extended N-terminal domains predicted to undergo liquid-liquid phase separation (Supplementary Figure 4e-g) or dimerize via α-helical coiled-coils (Supplementary Figure 4h-k). This variety in Ubl architecture suggests that ubiquitination by BilABCD operons could disrupt target-protein function through mislocalization, aggregation, or aberrant oligomerization.

In our structures of the E1^BilD^:E2^BilB^:Ubl^BilA^ complex, Ubl^BilA^ is bound to E1^BilD^ equivalently to known Ubl-E1 complexes, with its C-terminus positioned in the E1^BilD^ adenylation active site. We do not observe electron density for a bound ATP or AMP molecule in the active site. Indeed, the C-terminal alanine residue of Ubl^BilA^ (Ala98) is positioned in a manner that would physically prevent binding of ATP. In a sequence alignment of all Ubl^BilA^ homologs from Type II BilABCD operons, we found that these proteins encode a universally-conserved glycine followed by up to nine residues of non-conserved sequence (Supplementary Figure 4b). In *E. aridi* Ubl^BilA^, the conserved glycine (Gly97) is positioned one residue from the C-terminus. Close inspection of our structures reveals that Ubl^BilA^ residue Gly97 is positioned properly for adenylation, but cannot undergo this reaction because of the presence of the C-terminal residue Ala98 (Supplementary Figure 3c, g). Thus, in this system Ubl^BilA^ likely requires proteolytic processing by its cognate DUB^BilC^ to expose the reactive Gly97 residue for catalysis by E1^BilD^ (see next section).

In our structures, E2^BilB^ adopts an E2-like fold with the closest structural homologs being *S. cerevisiae* UBC12 (Cα r.m.s.d. 3.0 Å over 94 residue pairs)^30^ and several E2 proteins from *H. sapiens* including UBE2D2 (Cα r.m.s.d. 3.2 Å over 99 residues)^32^, UBE2D3 (Cα r.m.s.d. 3.4 Å over 96 residues)^33^, and UBE2J2 (Cα r.m.s.d. 3.4 Å over 93 residues)^34^ (Figure 2d). The conserved CEHH motif in this protein includes residues Cys138, Glu144, His146, and His151 ^8^. Of these, Cys138 is positioned equivalently to canonical E2 proteins’ catalytic cysteine. His151 is adjacent to Cys138, and may participate directly in catalysis. The other two conserved residues (Glu144 and His146) appear to play purely structural roles in E2^BilB^, with their side chains hydrogen-bonding to nearby backbone amide and carbonyl groups (Supplementary Figure 5a). In keeping with their likely functional importance, the putative catalytic cysteine (Cys138) and its adjacent histidine (His151) are highly conserved in E2^BilB^ proteins across both Type I and Type II BilABCD operons, while Glu144 and His146 are not conserved in Type I BilABCD (Supplementary Figure 5a). In both structures of the E1^BilD^:E2^BilB^:Ubl^BilA^ complex, E2^BilB^ is bound to the E1^BilD^ AAD near the adenylation active site and CYS domain (Figure 1b). While E1^BilD^ lacks the C-terminal UFD that participates in E2 binding in many canonical E1 proteins, the binding of E2^BilB^ to E1^BilD^ nonetheless closely resembles canonical E1-E2 binding. The interaction involves a loop and α-helix near the C-terminus of E1^BilD^ (residues 490-504), which is rigidified by the bound Zn^2+^ ion coordinated by Cys340 and Cys343 in the AAD crossover loop, plus Cys491 and Cys493 near the E1^BilD^ C-terminus (Figure 1d, Supplementary Figure 3d,h). In the E1^BilD^:E2^BilB^:Ubl^BilA^ Form 2 structure, the E1^BilD^ CYS domain is rotated away from the adenylation active site, and its putative catalytic cysteine is positioned adjacent to the E2^BilB^ putative catalytic cysteine (Cys138). In the structure, the two residues are apparently linked via a disulfide bond. This state closely mimics the structural state of an E1-to-E2 transthioesterification intermediate, in which the E1 and E2 catalytic cysteine residues approach to within ∼2 Å of one another^35^. Thus, our two structures reveal how the mobile E1^BilD^ CYS domain likely reacts first with a Ubl^BilA^-adenylate intermediate to generate the E1^BilD^∼Ubl^BilA^ thioester, then rotates upward to mediate Ubl^BilA^ handoff to E2^BilB^.

E2^BilB^ contains an extended disordered loop between strand β4 and the putative catalytic cysteine residue (Cys138), which is not found in other E2 proteins. This loop shows high sequence divergence among E2^BilB^ orthologs, and we propose that in the absence of an E3 substrate adapter protein in this pathway, this extended loop in E2^BilB^ may mediate specific target recognition. In both structures of the E1^BilD^:E2^BilB^:Ubl^BilA^ complex, E2^BilB^ forms a symmetric homodimer either through non-crystallographic symmetry (Form 1) or crystallographic symmetry (Form 2; Supplementary Figure 5b-d). Supporting the biological relevance of the observed E2^BilB^ dimer, an AlphaFold structure prediction with two copies of E2^BilB^ shows a nearly identical dimer architecture (Supplementary Figure 5e-f). Further supporting the idea that E2^BilB^ dimerizes in solution, the calculated molecular weight and hydrodynamic radius of a 2:2:2 E1^BilD^:E2^BilB^:Ubl^BilA^ complex closely matches the value we observed by size exclusion chromatography (Supplementary Figure 2). Homodimerization of E2 proteins has not been previously observed, and we theorize that E2^BilB^ dimerization may be functionally important for target-protein recognition.

### A bacterial DUB pre-processes its cognate Ubl

In eukaryotic Ubl pathways, deubiquitinases play two roles: they pre-process Ubls to expose their reactive C-terminal glycine (peptide bond cleavage), and they also cleave Ubl-target conjugates (isopeptide bond cleavage)^7^. Our structures and sequence alignments suggest that in many BilABCD systems including that of *E. aridi*, DUB^BilC^ is required to pre-process Ubl^BilA^ to expose the reactive C-terminal glycine. To test the activity of *E. aridi* DUB^BilC^, we generated a model substrate comprising Ubl^BilA^ fused at its C-terminus to green fluorescent protein (GFP). We found that wild type DUB^BilC^, but not two different active-site mutants (Glu33 to Ala (E33A) and Asp106 to Ala(D106A)), showed robust cleavage of Ubl^BilA^-GFP (Figure 3a). We isolated the C-terminal product of Ubl^BilA^-GFP cleavage and subjected it to N-terminal sequencing by Edman degradation, and found that DUB^BilC^ cleaves Ubl^BilA^ one residue upstream of the C-terminus, between Gly97 and Ala98 (Figure 3b, Supplementary Figure 6a). Thus, *E. aridi* DUB^BilC^ likely pre-processes Ubl^BilA^ to expose the reactive Gly97 for catalysis.

**Figure 3.**
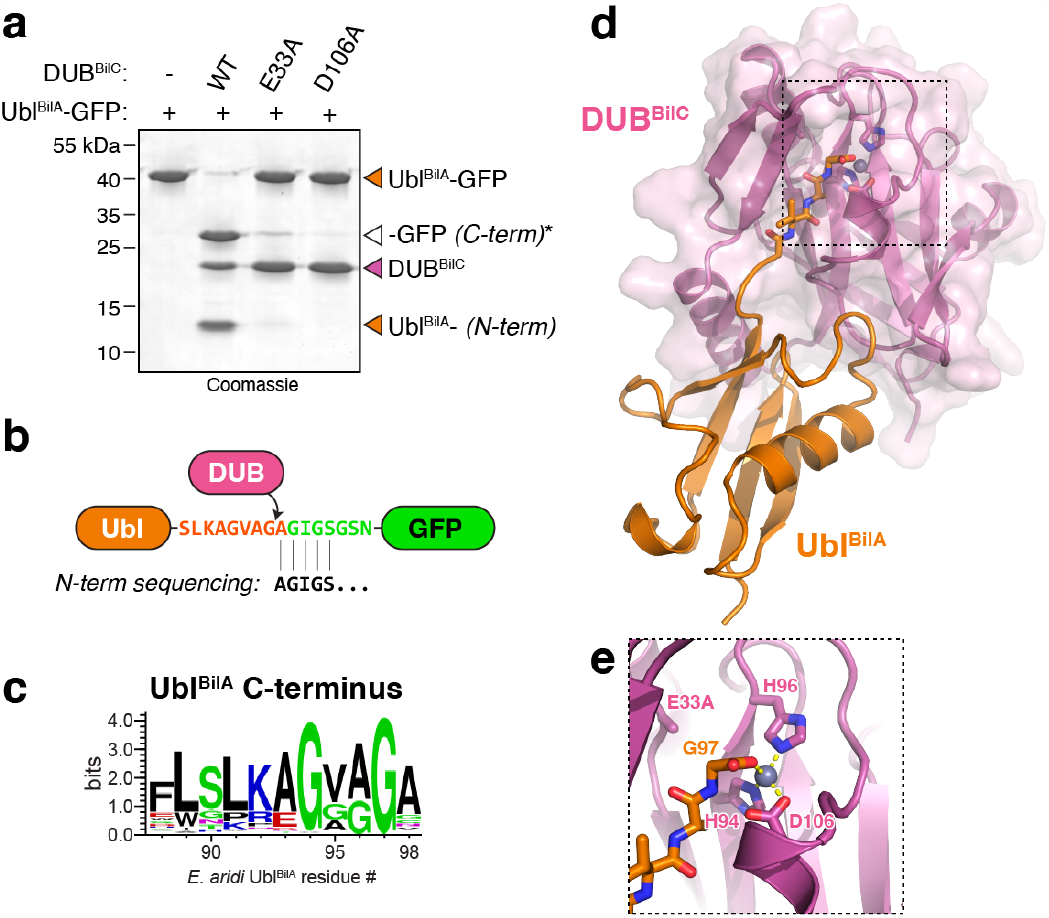
*E. aridi* DUB^BilC^ specifically cleaves Ubl^BilA.^ **(a)** Biochemical analysis of *E. aridi* DUB^BilC^ cleaving a Ubl^BilA^-GFP fusion protein. DUB^BilC^ constructs used were wild-type (WT), E33A active site mutant, or D106A active site mutant. **(b)** Schematic of the cleavage reaction shown in panel (a) and analysis by N-terminal sequencing (Edman degradation) of the C-terminal fragment (marked with an asterisk in panel (a)). See **Supplementary Figure 6a** for N-terminal sequencing data showing the sequence AGIGS, pinpointing the DUB^BilC^ cleavage site as C-terminal to Ubl^BilA^ Gly97. **(c)** Sequence logo from bacterial Type II Ubl^BilA^ proteins (Supplementary Table 1). Type II Ubl^BilA^ homologs possess up to nine residues C-terminal to the highly conserved glycine (Gly97 in *E. aridi* Ubl^BilA^). **(d)** X-ray crystal structure of DUB^BilC^(E33A) (purple) bound to Ubl^BilA^ (orange). A composite omit map (**Supplementary Figure 6b**) shows that Ubl^BilA^ is cleaved at Gly97. See Supplementary Figure 6d for structural comparisons with eukaryotic JAMM-family peptidases. **(e)** Closeup view of the DUB^BilC^(E33A) active site with bound zinc ion (gray) and the Ubl^BilA^ C-terminus (orange).

We next purified a complex of *E. aridi* DUB^BilC^(E33A) and full-length Ubl^BilA^, and determined two X-ray crystal structures at 1.36 Å resolution (Form 1) and 1.68 Å resolution (Form 2) (Figure 2c-d, Supplementary Figure 6b-c, Supplementary Table 2). The structures reveal DUB^BilC^ as a JAB/JAMM family peptidase, with the closest structural homologs being an archaeal JAMM peptidase and several eukaryotic AMSH enzymes (Supplementary Figure 6d). In both structures, Ubl^BilA^ is bound to DUB^BilC^ with its C-terminus positioned in the active site near the catalytic Zn^2+^ ion (Figure 3d-e). Despite using the catalytic mutant DUB^BilC^(E33A) for crystallization, we observed that in both structures Ubl^BilA^ was cleaved at residue Gly97. This observation is likely explained by residual low-level cleavage activity in this mutant (Figure 3a). These structural data support a model in which *E. aridi* DUB^BilC^ pre-processes Ubl^BilA^ by cleaving between Gly97 and Ala98.

### BilABCD is a fully functional ubiquitination system

Our structural data on E1^BilD^, E2^BilB^, and Ubl^BilA^ suggest that these proteins comprise a complete ubiquitination system equivalent to canonical eukaryotic ubiquitination pathways. To test this idea, we coexpressed E1^BilD^ and E2^BilB^ with Ubl^BilA^ truncated at Gly97 (Ubl^BilA^Δ97) in *E. coli*. Using an N-terminal His_6_-tag on Ubl^BilA^, we purified Ubl^BilA^ and associated proteins. In native buffer conditions that maintain both covalent and noncovalent complexes with Ubl^BilA^, we observed three major bands representing E1^BilD^, E2^BilB^, and Ubl^BilA^, plus minor bands with a wide range of molecular weights (Figure 4a). These minor bands were not present when Ubl^BilA^ contained its native C-terminal Ala98 residue; when E1 was mutated to eliminate Ubl^BilA^ adenylation (Arg246 to Ala); when the E1^BilD^ putative catalytic cysteine (Cys417) was mutated to alanine; or when the E2^BilB^ putative catalytic cysteine (Cys138) was mutated to alanine (Figure 4a). These data suggest that the observed minor bands represent covalent Ubl^BilA^-target protein conjugates that depend on Ubl^BilA^ adenylation by E1^BilD^, followed by sequential thioester formation with E1^BilD^ and E2^BilB^.

**Figure 4.**
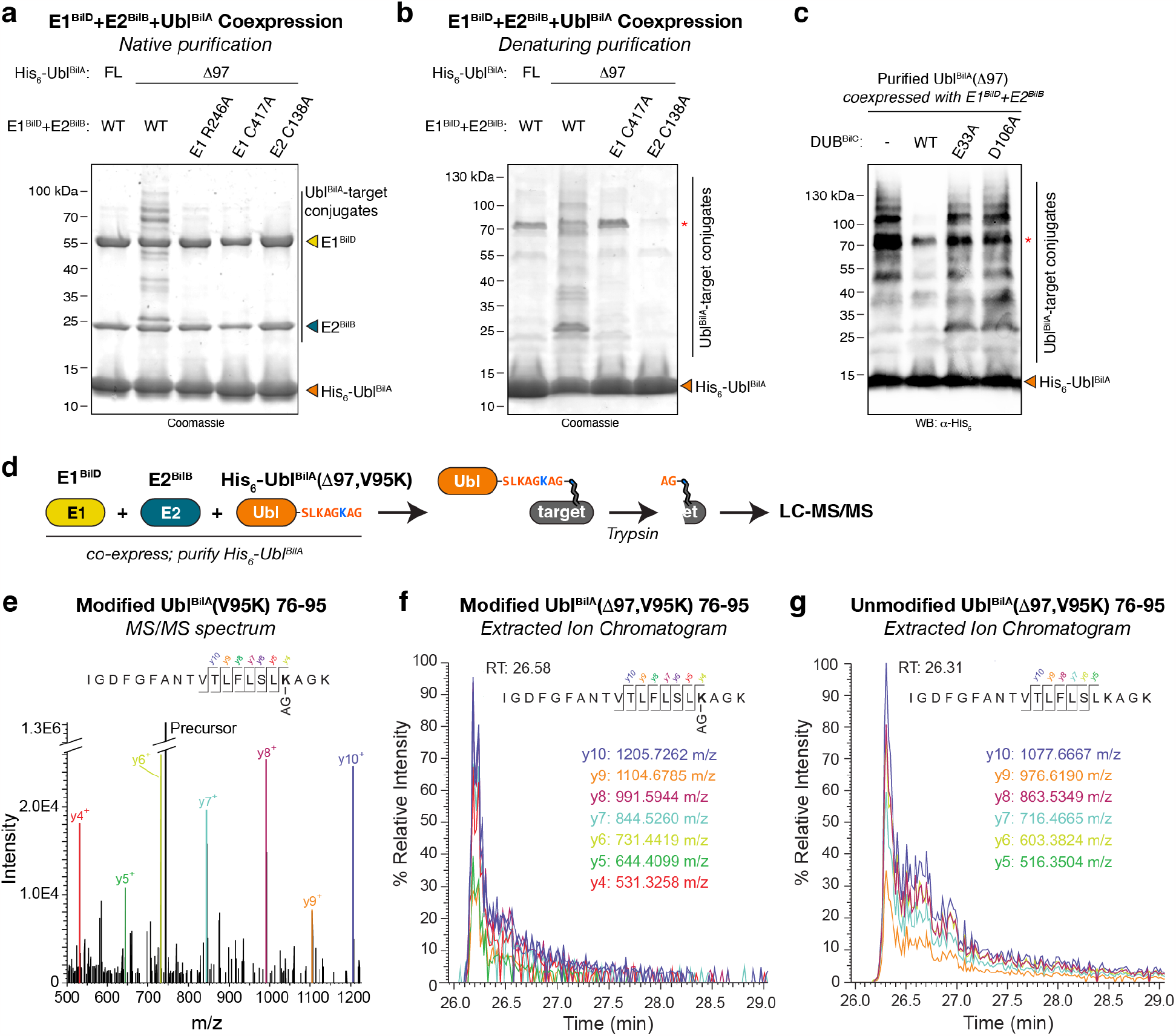
Bacterial BilABCD systems catalyze ubiquitination. **(a)** SDS-PAGE analysis of Ni^2+^ affinity-purified Ubl^BilA^ and associated proteins in native conditions, after coexpression with E1^BilD^ and E2^BilB^. Bands representing His_6_-Ubl^BilA^, E1^BilD^, and E2^BilB^ are marked. Ubl^BilA^-target conjugates are visible in the second lane. Ubl^BilA^ FL: full-length, unreactive Ubl^BilA^. Ubl^BilA^ Δ97: truncated at glycine 97 to enable E1-mediated catalysis; E1 R246A: adenylation active site mutant; E1 C417A: catalytic cysteine mutant; E2 C138A: catalytic cysteine mutant. WT: wild type. **(b)** SDS-PAGE analysis of Ni^2+^ affinity-purified Ubl^BilA^ and associated proteins in denaturing conditions, after coexpression with E1^BilD^ and E2^BilB^. Bands representing His_6_-Ubl^BilA^ and His_6_-Ubl^BilA^-target conjugates are marked. Red asterisk: contaminant. **(c)** SDS-PAGE and anti His_6_-tag western blot analysis of affinity-purified His_6_-Ubl^BilA^-target conjugates (from denaturing purifications) treated with DUB^BilC^. **(d)** Experimental scheme for identification of Ubl^BilA^ targets. See **Supplementary Figure 7a** for purification of His_6_-Ubl^BilA^(Δ97, V95K)-target complexes. **(e)** MS/MS spectrum of modified Ubl^BilA^(Δ97, V95K) residues 76-95. Y-ions shifted by the dipeptide Ala-Gly (AG) remnant are annotated in color. **(f)** Extracted Ion Chromatograms (EICs) of transition ions of the mis-cleaved, modified Ubl^BilA^(Δ97, V95K) residues 76-95, using a 10 ppm m/z tolerance. RT: retention time. **(g)** Extracted Ion Chromatograms (EICs) of transition ions of the mis-cleaved, unmodified Ubl^BilA^(Δ97, V95K) residues 76-95, using a 10 ppm m/z tolerance. RT: retention time. See **Supplementary Figure 7b** for EIC of the properly cleaved, unmodified peptide spanning Ubl^BilA^(Δ97, V95K) residues 76-92.

We next purified His_6_-tagged Ubl^BilA^Δ97 in denaturing conditions to eliminate copurification of non-covalently associated proteins. When coexpressed with wild-type E1^BilD^ and E2^BilB^, we again observed that Ubl^BilA^ copurified with a range of minor bands, which were not present when either the E1 or E2 putative catalytic cysteine residues were mutated to alanine (Figure 4b). These data confirm that the minor bands are likely covalent complexes between Ubl^BilA^ and *E. coli* proteins, that are generated by E1^BilD^ and E2^BilB^. We incubated the purified mixture of Ubl^BilA^-target conjugates with purified DUB^BilC^, then visualized the reactions by SDS-PAGE and western blotting with an antibody that recognizes the His_6_-tag on Ubl^BilA^. Untreated samples showed a large number of bands, suggesting that each band represents an *E. coli* protein conjugated to His_6_-tagged Ubl^BilA^ (Figure 4c). After incubation with wild-type DUB^BilC^, the minor bands were mostly lost and western blotting showed a single major band corresponding to monomeric Ubl^BilA^ (Figure 4c). The minor bands were not lost upon incubation with the DUB^BilC^ catalytic mutants E33A or D106A, confirming that this loss is due to proteolysis by DUB^BilC^. Thus, E1^BilD^ and E2^BilB^ can generate covalent Ubl^BilA^-target conjugates, which DUB^BilC^ can cleave.

Finally, we used mass spectrometry (MS) to verify that Ubl^BilA^ is conjugated to target proteins through an isopeptide linkage. We generated a His_6_-tagged Ubl^BilA^Δ97 construct with a Val95 to Lys (V95K) mutation, such that trypsin digestion of Ubl^BilA^-target conjugates would leave an alanine-glycine dipeptide linked to a target lysine residue (Figure 4d). We verified that the V95K mutant did not disrupt Ubl^BilA^-target conjugate formation (Supplementary Figure 7a), then purified His_6_-Ubl^BilA^(Δ97, V95K) in denaturing conditions after coexpression with E1^BilD^ and E2^BilB^. We separated the purified proteins by SDS-PAGE, then excised a region of the gel containing ∼20-40 kDa proteins (importantly, excluding monomeric Ubl^BilA^). We extracted proteins from the gel, cleaved with trypsin, and performed LC-MS/MS with data-dependent acquisition to search for peptides that contained a lysine residue modified with an alanine-glycine dipeptide. We identified a peptide corresponding to residues 76-95 of Ubl^BilA^(Δ97, V95K) modified at Lys92, then modified the MS parameters to selectively detect its daughter ions (Figure 4e). All detected daughter ions showed identical chromatographic elution profiles by reverse-phage liquid chromatography (Figure 4f), supporting the identification of this modified peptide. We similarly identified the unmodified Ubl^BilA^(Δ97, V95K) 76-95 peptide with an essentially identical chromatographic elution profile (Figure 4g), and the Ubl^BilA^(Δ97, V95K) 76-92 peptide that is properly cleaved by trypsin at Lys92 (Supplementary Figure 7b). These data confirm that E1^BilD^ and E2^BilB^ can catalyze formation of a *bona fide* Ubl^BilA^-target isopeptide linkage.

## Discussion

The evolutionary origins of ubiquitination and related pathways have remained mysterious since their discovery in the 1970s. While bacteria encode widespread E1-and Ubl-like proteins involved in sulfur metabolism, true ubiquitin-like protein conjugation was not demonstrated in bacteria until the CBASS-associated protein Cap2 was shown to conjugate a cGAS-like protein to target proteins as part of an antiviral immune response^17,18^. Here, we reveal that a different bacterial immune system, BilABCD, encodes a fully functional ubiquitination system with E1, E2, DUB, and Ubl proteins that architecturally and mechanistically resemble canonical eukaryotic ubiquitination machinery. We demonstrate that the bacterial E1 protein BilD encodes tandem IAD and AAD domains, with a mobile CYS domain capable of shuttling the Ubl protein BilA from the E1^BilD^ adenylation site to the E2 protein BilB. We show that all BilABCD proteins more closely resemble canonical eukaryotic ubiquitination machinery than functionally related bacterial proteins. When coexpressed in *E. coli*, E1^BilD^ and E2^BilB^ mediate Ubl^BilA^ conjugation to both cellular proteins and Ubl^BilA^ itself. Finally, the DUB protein BilC pre-processes Ubl^BilA^ to expose its reactive C-terminal glycine, and can also cleave Ubl^BilA^-target conjugates. Overall, these data show that bacteria encode ancient relatives of canonical ubiquitination systems, and suggest that ubiquitination-like protein conjugation evolved in the context of bacterial innate immunity.

Two bacterial immune systems are now known to catalyze ubiquitination-like protein conjugation: Type II CBASS systems encode Cap2, a combined E1-E2 protein that resembles the eukaryotic non-canonical E1 Atg7 and its partner E2s Atg3 and Atg10 ^17^, while BilABCD pathways’ Ubl^BilA^, E1^BilD^, and E2^BilB^ resemble canonical eukaryotic Ubl, E1, and E2 proteins. Remarkably, several other uncharacterized bacterial operon families are predicted to encode distinct combinations of Ubl, E1, E2, RING E3, and DUB proteins^8-11,17^. As with Type II CBASS and BilABCD systems, these operon families are predicted to function in antiviral defense^8-11^. Thus, the evolutionary pressure of the phage-bacteria arms race has given rise to a profusion of ubiquitination-like protein conjugation pathways that likely interrupt phage infections through a variety of mechanisms. Our data reveal that when expressed in a heterologous host, *E. aridi* E1^BilD^ and E2^BilB^ can mediate nonspecific Ubl^BilA^-target conjugation. In the context of bacteriophage infection, BilABCD could protect its host by non-specifically ubiquitinating abundant phage proteins, potentially inhibiting their self-assembly into functional phage progeny. A more likely scenario is that BilABCD specifically ubiquitinates one or a few phage proteins to interrupt key life-cycle events like DNA replication, virion assembly, or host-cell lysis. Alternatively, BilABCD could cause production of progeny phage that are non-infectious due to ubiquitination of key structural proteins. Indeed, recent work on a Type I BilABCD operon from *Collimonas sp*. OK412 shows that this system specifically ubiquitinates the central tail fiber protein of phages Secphi27 and Secphi4, and that this activity results in both defective virion assembly and impaired infectivity of progeny phage^14^. We propose that the extended disordered loop in E2^BilB^ may recognize phage target proteins, providing specificity to this system in the absence of a dedicated E3 protein.

BilABCD operons were originally dubbed “<u>B</u>acterial ISG15-like” because of shared innate-immune function and structural similarity between Ubl^BilA^ and the eukaryotic Ubl ISG15 ^13^. Our data solidify the functional parallels between bacterial BilABCD pathways and eukaryotic innate-immune pathways that involve Ubl conjugation, including the ISG15 pathway. In bacterial immunity, Ubl conjugation in different BilABCD systems likely has diverse effects on infecting phages, including the known effects on virion assembly and infectivity through modification of a key tail fiber protein^14^, plus potentially aberrant oligomerization or aggregation of modified target proteins mediated by diverse Ubls. The ISG15 pathway has been proposed to function similarly, by modifying viral proteins to interrupt key life-cycle events^3,36,37^. While the ISG15 pathway appears to have emerged after the establishment of eukaryotes rather than being inherited from a bacterial or archaeal ancestor^9^, the functional parallels between BilABCD and ISG15 pathways strongly point to the effectiveness of ubiquitination-like protein conjugation pathways in mediating innate immunity across kingdoms.

## Acknowledgements

The authors thank the staff at The Protein Facility of the Iowa State University Office of Biotechnology for assistance with Edman degradation, R. Sorek for sharing information prior to publication, and members of the Corbett laboratory for helpful discussions. This work was funded by NIH R35 GM144121 (K.D.C.); the NIH Director’s New Innovator Award DP2 AT012346 (A.T.W.), a Mallinckrodt Foundation Grant (A.T.W.), the Boettcher Foundation’s Webb-Waring Biomedical Research Program (A.T.W.), and the Pew Biomedical Scholars Program (A.T.W.); NIH R01 GM116897 (H.Z. and R.T.S.), R01 GM151191 (H.Z. and R.T.S.), and S10 OD023498 (H.Z.). L.R.C. is supported by the UCSD Molecular Biophysics Training Grant (T2 GM139795), J.C. is supported by a Pfizer-Cell Signaling San Diego graduate fellowship, and H.E.L. is supported as a fellow of the Jane Coffin Childs Memorial Fund for Medical Research. This work is based upon research conducted at the Northeastern Collaborative Access Team beamlines, which are funded by the National Institute of General Medical Sciences from the National Institutes of Health (P30 GM124165). The Eiger 16M detector on 24-ID-E is funded by a NIH-ORIP HEI grant (S10OD021527). This research used resources of the Advanced Photon Source, a U.S. Department of Energy (DOE) Office of Science User Facility operated for the DOE Office of Science by Argonne National Laboratory under Contract No. DE-AC02-06CH11357. This paper was typeset with the bioRxiv word template by @Chrelli: www.github.com/chrelli/bioRxiv-word-template.

## Author contributions

L.R.C. designed experiments, cloned expression vectors, purified proteins, determined crystal structures of E1^BilD^:E2^BilB^:Ubl^BilA^, performed E1^BilD^:E2^BilB^:Ubl^BilA^ coexpression/activity assays, and wrote the paper with Q.Y. and K.D.C.. Q.Y. designed experiments, cloned expression vectors, purified proteins, determined crystal structures of DUB^BilC^:Ubl^BilA^ and *M. brachiatum* Ubl^BilA^, performed DUB^BilC^ activity assays, and wrote the paper with L.R.C. and K.D.C.. J.C. designed experiments and performed mass spectrometry analysis of Ubl^BilA^-associated proteins. M.G. cloned expression vectors, purified proteins, and assisted with crystal structures of DUB^BilC^:Ubl^BilA^. H.E.L. performed bioinformatics analysis. H.Z. oversaw mass spectrometry analysis. A.T.W. designed experiments and oversaw bioinformatics analysis. R.T.S. designed experiments and oversaw mass spectrometry analysis. K.D.C. designed experiments, oversaw structural and biochemical assays, and wrote the paper with L.R.C. and Q.Y..

## Materials and Methods

### Bioinformatics

To identify bacterial CEHH operons, the *Ensifer aridi* TW10 E1^BilD^ protein (IMG accession 2509379665) was used as a query for BLAST searches in the IMG database of bacterial genomes (https://img.jgi.doe.gov/). Hit sequences were filtered for redundancy, aligned using the MAFFT^38^ version 7 web server (https://mafft.cbrc.jp/alignment/server/index.html) and visualized in Jalview^39^. For each non-redundant hit, the genomic neighborhood of the E1^BilD^ gene was visually inspected in IMG for neighboring genes similar to *E. aridi* Ubl^BilA^, E2^BilB^, and DUB^BilC^, plus genes similar to CapH (usually annotated as Pfam01381; XRE-family HTH domain) and CapP (usually annotated as Pfam06114; Peptidase_M78)^23^. In all cases, the *bilA-bilB-bilC-bilD* gene order was consistent, and in all but one case *capH* and *capP* were positioned upstream of *bilABCD* and oriented in the opposite coding direction (i.e. sharing a common promoter region with *bilABCD*). Average linkage (UPGMA) trees were calculated by the MAFFT web server and visualized using the Interactive Tree of Life server (https://itol.embl.de/). Protein structure predictions (monomer and multimer) were performed using the AlphaFold_MMseqs2 implementation of ColabFold^40^, which implements AlphaFold 2 ^41,42^ predictions using sequence alignments generated by MMseqs2 ^43^.

### Protein expression and purification

*Ensifer aridi* Ubl^BilA^ (Joint Genome Institute Integrated Microbial Genomes (JGI-IMG) accession 2509379668), E2^BilB^ (IMG accession 2509379667), DUB^BilC^ (IMG accession 2509379666), and E1^BilD^ (IMG accession 2509379665), plus *Methylobacterium brachiatum* DSM 19569 Ubl^BilA^ (IMG accession 2928979547) were individually cloned into *E. coli* expression vectors encoding either no tag (UC Berkeley Macrolab vector 2A-T, Addgene ID 29665) or an N-terminal TEV protease-cleavable His_6_-tag (UC Berkeley Macrolab vector 2B-T, Addgene ID 29666). For coexpression, multigene coexpression cassettes were generated by PCR and cloned into UC Berkeley Macrolab vector 2B-T to generate an N-terminal TEV protease-cleavable His_6_-tag on one protein. Targeted mutants were generated by PCR-based mutagenesis.

Vectors were transformed into *E. coli* Rosetta 2 pLysS (EMD Millipore), and 1 L cultures were grown at 37°C in 2XYT media to an OD_600_ of 0.6 before induction with 0.25 mM IPTG at 20°C for 16-18 hours. Cells were harvested by centrifugation, resuspended in buffer A (25 mM Tris-HCl pH 8.5, 300 mM NaCl, 5 mM MgCl_2_, 10% glycerol and 5 mM mercaptoethanol) containing 5 mM imidazole, then lysed by sonication (Branson Sonifier). Lysates were clarified by centrifugation, then supernatants were passed over a Ni-NTA Superflow column (Qiagen) in resuspension buffer. The column was washed in wash buffer (buffer A containing 20 mM imidazole), then eluted in elution buffer (buffer A containing 400 mM imidazole). Eluates were concentrated by ultrafiltration (Amicon Ultra; EMD Millipore), then passed over a Superdex 200 Increase size exclusion column (Cytiva) in size exclusion buffer (25 mM Tris-HCl pH 8.5,300 mM NaCl, 5 mM MgCl_2_, 10% glycerol and 1mM DTT). Peak fractions were concentrated by ultrafiltration and stored at 4°C.

For denatured purification of E1^BilD^:E2^BilB^:His_6_-Ubl^BilA^, cells were harvested by centrifugation, resuspended in buffer A, then lysed by sonication (Branson Sonifier). Lysates were clarified by centrifugation, then supernatants were passed over a Ni-NTA Superflow column (Qiagen) in resuspension buffer. The column was washed in buffer B (100 mM Tris-HCl pH 8, 8 M Urea and 100 mM NaH_2_PO_4_), followed by wash buffer (buffer A containing 20 mM imidazole), then eluted in elution buffer (buffer A containing 400 mM imidazole). Eluates were concentrated by ultrafiltration (Amicon Ultra; EMD Millipore) and stored at 4°C.

### DUB^BilC^ activity assays and N-terminal sequencing

To generate a model substrate for DUB^BilC^ cleavage, Ubl^BilA^ was cloned into a vector encoding a C-terminal GFP tag (UC Berkeley Macrolab vector H6-msfGFP, Addgene ID 29725) and purified as above. The Ubl^BilA^-GFP fusion protein (10 μg) was mixed with 5 μg of DUB^BilC^ (wild type or mutants) in 20 μL reaction buffer containing 20 mM HEPES pH 7.5, 100 mM NaCl, 20 mM MgCl_2_, 20 μM ZnCl_2_ and 1 mM DTT, then incubated 30 minutes at 37°C. Reactions were analyzed by SDS-PAGE with Coomassie blue staining. For protein N-terminal sequencing by Edman degradation, cleavage products were separated by SDS-PAGE, transferred to a Bio-Rad ImmunBlot PVDF membrane, and visualized by staining with Coomassie Blue R-250. The band representing the C-terminal cleavage product of Ubl^BilA^-GFP was excised, the membrane was washed with methanol and deionized water, then loaded onto a Shimadzu PPSQ-53A instrument for analysis. Five rounds of Edman degradation were performed, and the “evaluated value” scores for each round were analyzed to obtain the likely N-terminal sequence.

For western blots of His_6_-Ubl^BilA^ conjugates, proteins were transferred to PVDF membranes using a Trans-Blot Turbo RTA Mini 0.2 μm PVDF Transfer Kit (Bio-Rad) according to the manufacturer’s instructions, using a Trans-Blot Turbo Transfer System (Bio-Rad). Membranes were blocked with 5% Non-Fat Dry Milk in TBST, then incubated with mouse anti-His tag primary antibody (Millipore Sigma SAB1305538) at 1:1,000 dilution, followed by HRP-conjugated secondary antibody (HRP Goat anti-mouse IgG, Millipore Sigma AP128P) at 1:30,000 dilution. After washing, HRP signal was detected with Amersham ECL Select Western Blotting Detection Reagent (Cytiva) using a ChemiDoc Imaging System (Bio-Rad) in Protein Blot -Chemiluminescence setting.

### Crystallography

For crystallization of the *E. aridi* DUB^BilC^(E33A):Ubl^BilA^ complex (Form 1), purified protein at 30 mg/mL in crystallization buffer (25 mM Tris-HCl pH 8.5, 200 mM NaCl, 5 mM MgCl_2_, and 1 mM TCEP (tris(2-carboxyethyl)phosphine)) was mixed 1:1 with well solution containing 100 mM HEPES pH 7.5, 0.2 M MgCl_2_, and 25% PEG 3350 in hanging drop format. Crystals were harvested into cryoprotectant solution containing an additional 10% glycerol and frozen in liquid nitrogen. Diffraction data were collected at the Advanced Photon Source (Argonne National Lab) NE-CAT beamline 24ID-E on April 7, 2023 (collection temperature 100 K; x-ray wavelength 0.97918 Å) (Supplementary Table 2). Data were processed with the RAPD data-processing pipeline (https://github.com/RAPD/RAPD), which uses XDS^44^ for data indexing and reduction, POINTLESS^45^ for space group assignment, and AIMLESS^46^ for scaling. The structure was determined by molecular replacement in PHASER^47^ using a predicted structure from AlphaFold 2 ^41^ as a search model. The model was manually rebuilt in COOT^48^ and refined in phenix.refine^49^ using positional and individual B-factor refinement. B-factors for all atoms except waters were refined anisotropically. The final model has good geometry, with 98.75% of residues in favored Ramachandran space, 1.25% allowed, and 0% outliers. The overall MolProbity score is 1.18, and the MolProbity clash score is 3.98.

For crystallization of the *E. aridi* DUB^BilC^(E33A):Ubl^BilA^ complex (Form 2), purified protein at 30 mg/mL in crystallization buffer (25 mM Tris-HCl pH 8.5, 200 mM NaCl, 5 mM MgCl_2_, and 1 mM TCEP (tris(2-carboxyethyl)phosphine)) was mixed 1:1 with well solution containing 100 mM MES pH 6.5 and 1M sodium/potassium tartrate. Crystals were harvested into cryoprotectant solution containing an additional 30% glycerol and frozen in liquid nitrogen. Diffraction data were collected at the Advanced Photon Source (Argonne National Lab) NE-CAT beamline 24ID-E on April 7, 2023 (collection temperature 100 K; x-ray wavelength 0.97918 Å) (Supplementary Table 2). Data were processed with the RAPD data-processing pipeline, and the structure was determined by molecular replacement in PHASER using the Form 1 BilC^E33A^:BilA structure (with ligands and waters excluded) as a search model. The model was manually rebuilt in COOT and refined in phenix.refine using positional and individual isotropic B-factor refinement. The final model has good geometry, with 99.58% of residues in favored Ramachandran space, 0.42% allowed, and 0% outliers. The overall MolProbity score is 1.08, and the MolProbity clash score is 1.86.

For crystallization of the *E. aridi* E1^BilD^:E2^BilB^:Ubl^BilA^ complex (Form 1), purified protein at 10 mg/mL in crystallization buffer was mixed 1:1 with well solution containing 100 mM HEPES pH 7.5, 100 mM sodium citrate, 5% isopropanol, and 10% PEG 3350 in hanging drop format. Crystals were harvested into cryoprotectant solution containing an additional 20% glycerol and frozen in liquid nitrogen. Diffraction data were collected at the Advanced Photon Source (Argonne National Lab) NE-CAT beamline 24ID-E on April 4, 2023 (collection temperature 100 K; x-ray wavelength 0.97918 Å) (Supplementary Table 2). Data were processed with the RAPD data-processing pipeline, and the structure was determined by molecular replacement in PHASER using predicted structures of each protein (E1^BilD^ CYS domain excluded) from AlphaFold2 as a search model. One copy of Ubl^BilA^, two copies of E2^BilB^, and two copies of E1^BilD^ were located. The model was manually rebuilt in COOT and refined in phenix.refine using positional and individual isotropic B-factor refinement. The final model has good geometry, with 96.23% of residues in favored Ramachandran space, 3.47% allowed, and 0.30% outliers. The overall MolProbity score is 1.64, and the MolProbity clash score is 5.86.

For crystallization of the *E. aridi* E1^BilD^:E2^BilB^:Ubl^BilA^ complex (Form 2), purified protein at 10 mg/mL in crystallization buffer was mixed 1:1 with well solution containing 100 mM imidazole pH 8.0, 200 mM MgCl_2_, and 10% PEG 3350 in hanging drop format. Crystals were harvested into cryoprotectant solution containing an additional 20% glycerol and frozen in liquid nitrogen. Diffraction data were collected at the Advanced Photon Source (Argonne National Lab) NE-CAT beamline 24ID-E on April 7, 2023 (collection temperature 100 K; x-ray wavelength 0.97918 Å) (Supplementary Table 2). Data were processed with the RAPD data-processing pipeline, and the structure was determined by molecular replacement in PHASER using predicted structures of each protein (E1^BilD^ CYS domain excluded) from AlphaFold2 as a search model. The model was manually rebuilt in COOT and refined in phenix.refine using positional and individual isotropic B-factor refinement. The final model has good geometry, with 96.05% of residues in favored Ramachandran space, 3.53% allowed, and 0.42% outliers. The overall MolProbity score is 1.82, and the MolProbity clash score is 5.62.

For crystallization of *M. brachiatum* Ubl^BilA^, purified protein was subjected to surface lysine methylation by treating with borane (50 mM final concentration) and formaldehyde (100 mM final concentration) for 1 hour at 4°C, followed by addition of glycine (25 mM final concentration) to quench the reaction for 30 minutes, followed by buffer exchange by centrifugal concentration. Methylated protein at 12 mg/mL in crystallization buffer was mixed 1:1 with well solution containing 100 mM CAPS pH 10.5, 0.2 M Li_2_SO_4_, 1.2 M NaH_2_PO_4_, and 0.8 M K_2_HPO_4_ in sitting drop format. Crystals were harvested into cryoprotectant solution containing 30% glycerol and frozen in liquid nitrogen. Diffraction data were collected at the Advanced Photon Source (Argonne National Lab) NE-CAT beamline 24ID-C on March 2, 2023 (collection temperature 100 K; x-ray wavelength 0.97911 Å) (Supplementary Table 2). Data were processed with the RAPD data-processing pipeline, and the structure was determined by molecular replacement in PHASER using a predicted structure from AlphaFold2 as a search model. Despite crystallizing full-length protein, only β-grasp domains 1 and 2 were visible in the model. The model containing four protomers was manually rebuilt in COOT and refined in phenix.refine using positional and individual isotropic B-factor refinement. The final model has good geometry, with 99.66% of residues in favored Ramachandran space, 0.34% allowed, and 0% outliers. The overall MolProbity score is 1.16, and the MolProbity clash score is 3.65.

Final coordinates and structure factors for all structures have been deposited at the RCSB Protein Data Bank (https://www.rcsb.org) under accession codes 8TYX (DUB^BilC^(E33A):Ubl^BilA^ Form 1), 8TYY (DUB^BilC^(E33A):Ubl^BilA^ Form 2), 8TZ0 (E1^BilD^:E2^BilB^:Ubl^BilA^ Form 1), 8TYZ (E1^BilD^:E2^BilB^:Ubl^BilA^ Form 2), and 8U38 (*M. brachiatum* Ubl^BilA^). Raw diffraction images have been deposited at the SBGrid Data Bank (https://data.sbgrid.org) under accession codes 1039 (DUB^BilC^(E33A):Ubl^BilA^ Form 1), 1040 (DUB^BilC^(E33A):Ubl^BilA^ Form 2), 1041 (E1^BilD^:E2^BilB^:Ubl^BilA^ Form 1), 1042 (E1^BilD^:E2^BilB^:Ubl^BilA^ Form 2), and 1043 (*M. brachiatum* Ubl^BilA^).

### In-Gel Digestion

For mass spectrometry identification of Ubl^BilA^-conjugated proteins, *E. aridi* E1^BilD^ and E2^BiLB^ were coexpressed with His_6_-Ubl^BilA^(V95K) in *E. coli*, and purified by Ni-NTA chromatography in denatured solution as above. Purified proteins were separated by SDS-PAGE and visualized with Coomassie blue staining, then mass spectrometry was performed by in-gel trypsin digestion as previously described^50^. In brief, gel slices were cut into ∼1 mm x 1 mm x 1 mm cubes using a clean razor blade in a glass dish. Gel slices were reduced in 100 μl of 10 mM DTT for 30 minutes at 37°C. Cysteine alkylation was performed by incubating the samples at room temperature in the dark for 20 minutes following the addition of 6 μl of 0.5 M iodoacetamide. To digest proteins, 100 μl of 10 ng/μl trypsin (Promega #V511A) in 20 mM ammonium bicarbonate (pH 8) was added to submerge the gel pieces, then incubated on ice for 30 minutes until fully swollen. An additional 20-50 μl of ammonium bicarbonate buffer was added to ensure the gel slices were fully submerged prior to overnight incubation at 37°C. The next day, trypsin digested peptides were extracted from the sample via multiple extractions using 50% acetonitrile/5% formic acid, dried under vacuum and reconstituted in 20 μl of 0.1% trifluoroacetic acid (pH 2).

### LC-MS/MS Analysis

Peptides were analyzed by LC-MS/MS on a Vanquish Neo high performance liquid chromatography system coupled to a Q-Exactive Plus mass spectrometer (Thermo Fisher Scientific). Data-dependent analysis (DDA) was performed using a Top 10 approach using a linear gradient of 2 – 30% mobile phase B (Mobile Phase A: 0.1% formic acid in H_2_O, Mobile Phase B: 0.1% formic acid in ACN) at a flow rate of 75 μL/min on a 15-cm C_18_ column maintained at 40°C (Acclaim Pepmap 1 mm I.D., 2 μm particle size, 100 Å pore size; Thermo Fisher Scientific). The LC method for DDA starts at 2% B and is held constant for 1 min, followed by a change to 30% B across 20 min, followed by a change to 70% B across 3 min, and then finally increased to 95% B across 0.1 min. Column washing and equilibration is performed across 4.1 min making the total method time 28.2 min. Parallel reaction monitoring (PRM) was performed in DIA mode utilizing a global inclusion list with a full scan using a linear gradient of 2 – 40% mobile phase B at a flow rate of 1.5 μL/min on an in-house packed 15-cm pulled-tip column emitter (100 μm I.D. with 2.2-μm C_18_ beads, 120 Å pore size; Sepax Technologies) maintained at 50 °C (PRSO-V2; Sonation GmbH). The LC method starts at 2% B and is held constant for 5 min, followed by a change to 40% B across 25 min, and then finally increased to 95% B across 1.5 min. Column washing and equilibration is performed across 13.5 min making the total method time 45 min. Targeted PRM acquisition of the Ubl^BilA^ (V95K) C-terminal peptides were performed in DIA mode using a targeted inclusion list for the +2 and +3 charge states for the modified and non-modified peptide species. Initial data review and quality control of PRM acquired transition spectra were analyzed in Skyline (v22.2.0.527). Final extracted transition ion chromatograms were generated using FreeStyle (v1.7, Thermo Fisher Scientific) using a ± 10 ppm m/z tolerance for each transition.

## Data-dependent Peptide Sequencing Analysis

Database searching of data-dependent acquired MS spectra was performed using the Trans-Proteomic Pipeline software suite v6.3.2 Arcus (TPP, Seattle Proteome Center)^51,52^. The search parameter file and protein database used for the database search is included in Supplementary Table 3. Briefly, database search was performed using COMET and peptides were quantified using XPRESS in label-free mode. A static modification of 57.021464 Da was applied to cysteine residues. Two differential modifications were applied as follows: Oxidation of methionine (15.9949 Da) and the expected AG remnant of Ubl^BilA^ (V95K) on lysine/N-termini (128.05857 Da). Quality control was performed following data analysis and manual inspection of chromatography and MS/MS spectral assignments. Redundant peptide identifications were removed to generate the final list of unique peptides (Supplementary Table 4).

The protein database included all proteins from the *E. coli* proteome (Uniprot: ECOLX) plus the following:

>tr|HIS6_Ubl-BilA_delta97-V95K

MKSSHHHHHHENLYFQSNASKDSRKGDNHGGGSGKIEIIVVVNGQPTQVEANPNQPLHVV

RTKALENTQNVAQPPDNWEFKDEAGNLLDVDKKIGDFGFANTVTLFLSLKAGKAG

>tr|E1-BilD

MALANFIDRAATAASQVLTDFHLGDFKAALEKQVVAVAFDDQAISCAEGQATLDLAVRLLA RLYPVLAILPLDSAASSQAQALERLAKSINRKIGIRRSGKSATVCLVAGATRPSLRCPTFF IGSDGWAAKLSRTDPVGSGSSLLPYGAGAASCFGAANVFRTIFAAQLTGAESDENIDLSLY SYNKSRAGDAGPIDPAVDLGETHLVGLGAIAHGALWALARQSGLSGRLHVVDHEAVELSNL QRYVLAGQAEIGMSKAVLATTALRSTALEVEAHPLKWAEHVARRGDWIFDRVGVALDTAAD RVAVQGALPRWIANAWTQEHDLGISRHGFDDGQACLCCMYMPSGKSKDEHQLVAEELGIPE AHEQVKALLQTNAGVPNDFVVRVATAMGVPFEPLAPFVGQPLRSFYQQAICGGLVFQLSDG SRLVRTVVPMAFQSALAGIMLAAELVKHSAGFPMSPTTSTRVNLLRPLGSHLHDPKAKDSSGRCICSDEDFISAYRRKYGNGVEPLSNISAEQKRTSPLPRTGRQVCA

>tr|E2-BilB

MPELQTVDPEVSRAKFDREISRFRPYADAYRMQGCFLIEESFPSAFFIFASPKVKPRVIGA AIEIDFTNYDLRPPSVVFVDPFTRQPIARKDLPLNMLRRPQLPGTPPEMISNLIQQNAVSLT DFIQANSLQDSPFLCMAGVREYHDNPAHSGDPWLLHRGSGEGCLAFILDKIIKYGTGPVEQL HIQLQYAVGLLVPPQAIPE

**Supplementary Figure 1.**
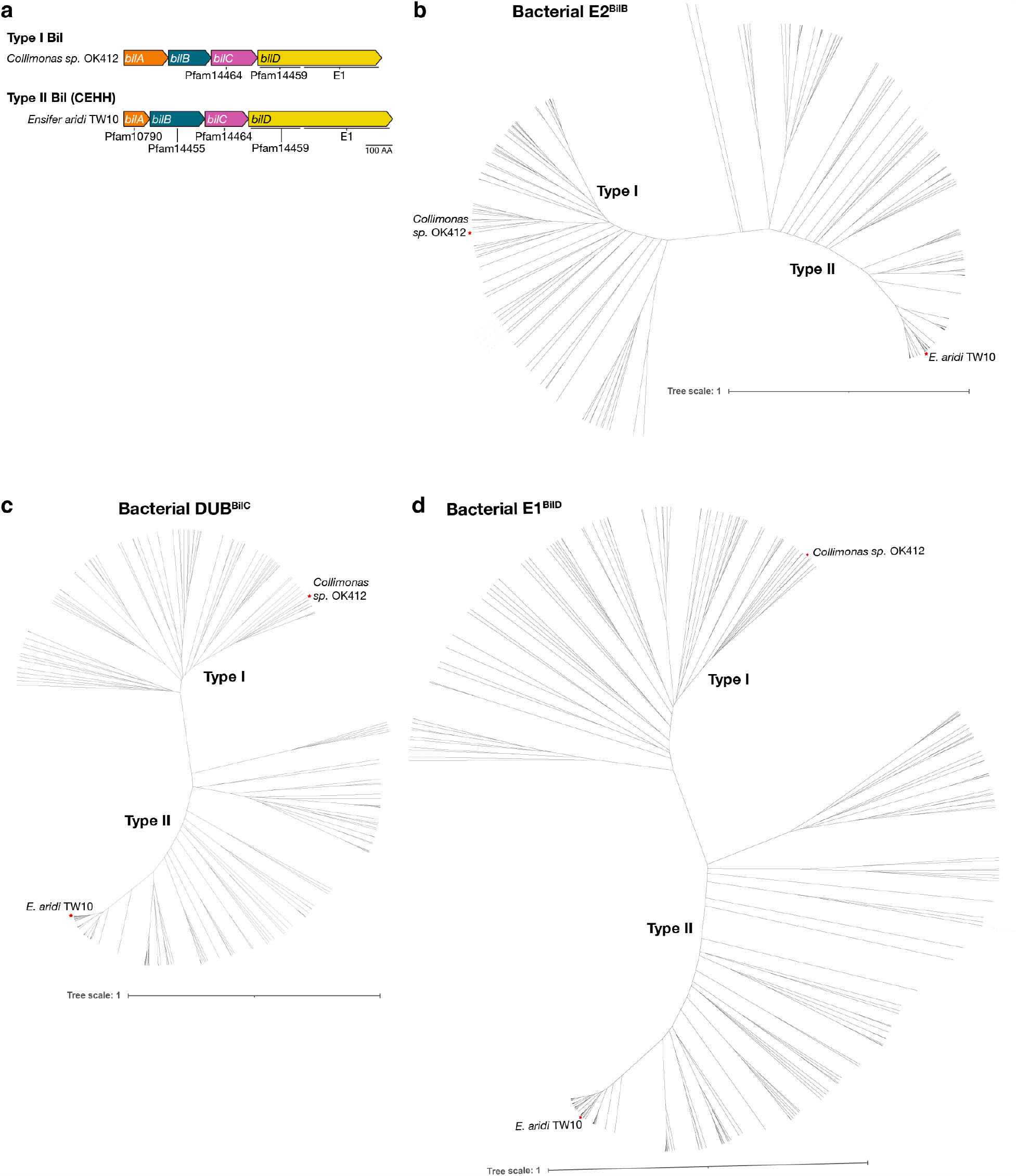
Comparison of Type I and Type II BilABCD operons. **(a)** Operon schematics of a Type I BilABCD operon from *Collimonas sp*. OK412 ^13^, and a Type II BilABCD operon from *E. aridi* TW10 (**Supplementary Table 1**). Noted under each gene are the conserved PFAM domain annotations for that gene. **(b)** Unrooted average distance tree assembled from E2^BilB^ proteins from Type I BilABCD operons^13^ and Type II BilABCD systems (**Supplementary Table 1**). Branches representing proteins from Type I and Type II operons are marked. Specific E2^BilB^ proteins from *Collimonas sp*. OK412 (Type I) and *E. aridi* TW10 (Type II) are indicated with red stars and labeled. Scale bar: 1 substitution per site. **(c)** Unrooted average distance tree assembled from DUB^BilC^ proteins from Type I BilABCD operons^13^ and Type II BilABCD operons (**Supplementary Table 1**). Scale bar: 1 substitution per site. Branches representing proteins from Type I and Type II operons are marked. Specific DUB^BilC^ proteins from *Collimonas sp*. OK412 (Type I) and *E. aridi* TW10 (Type II) are indicated with red stars and labeled. Scale bar: 1 substitution per site. **(d)** Unrooted average distance tree assembled from E1^BilD^ proteins from Type I BilABCD operons^13^ and Type II BilABCD operons (**Supplementary Table 1**). Scale bar: 1 substitution per site. Branches representing proteins from Type I and Type II operons are marked. Specific E1^BilD^ proteins from *Collimonas sp*. OK412 (Type I) and *E. aridi* TW10 (Type II) are indicated with red stars and labeled. Scale bar: 1 substitution per site.

**Supplementary Figure 2.**
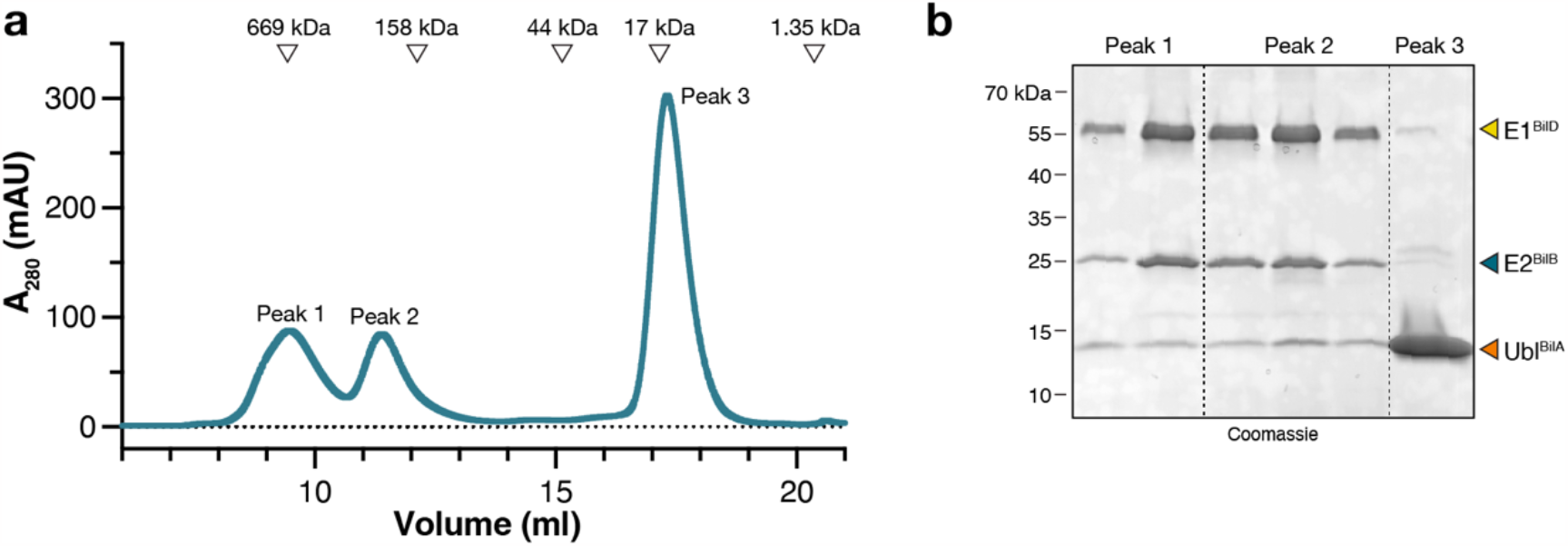
Purification and characterization of the *E. aridi* E1^BilD^:E2^BilB^:Ubl^BilA^ complex. **(a)** Size exclusion chromatography analysis of purified *E. aridi* E1^BilD^:E2^BilB^:Ubl^BilA^. Elution positions for molecular weight standards are shown at top. The apparent molecular weight and Stokes radius for Peak 2 are 249 kDa and 57.6 Å, respectively. The molecular weight of a 2:2:2 E1^BilD^:E2^BilB^:Ubl^BilA^ complex is 180.4 kDa, and the rotational hydrodynamic radius is 59.8 Å (as calculated by HullRad^54^ from the structure of a 2:2:2 complex built from the E1^BilD^:E2^BilB^:Ubl^BilA^ Form 2 crystal structure. Peak 1 likely represents aggregated E1^BilD^:E2^BilB^:Ubl^BilA^ complexes, and Peak 3 represents excess His_6_-tagged Ubl^BilA^. **(b)** SDSPAGE analysis of selected fractions from panel (a). Fractions from Peak 2 were concentrated and used for crystallization of E1^BilD^:E2^BilB^:Ubl^BilA^.

**Supplementary Figure 3.**
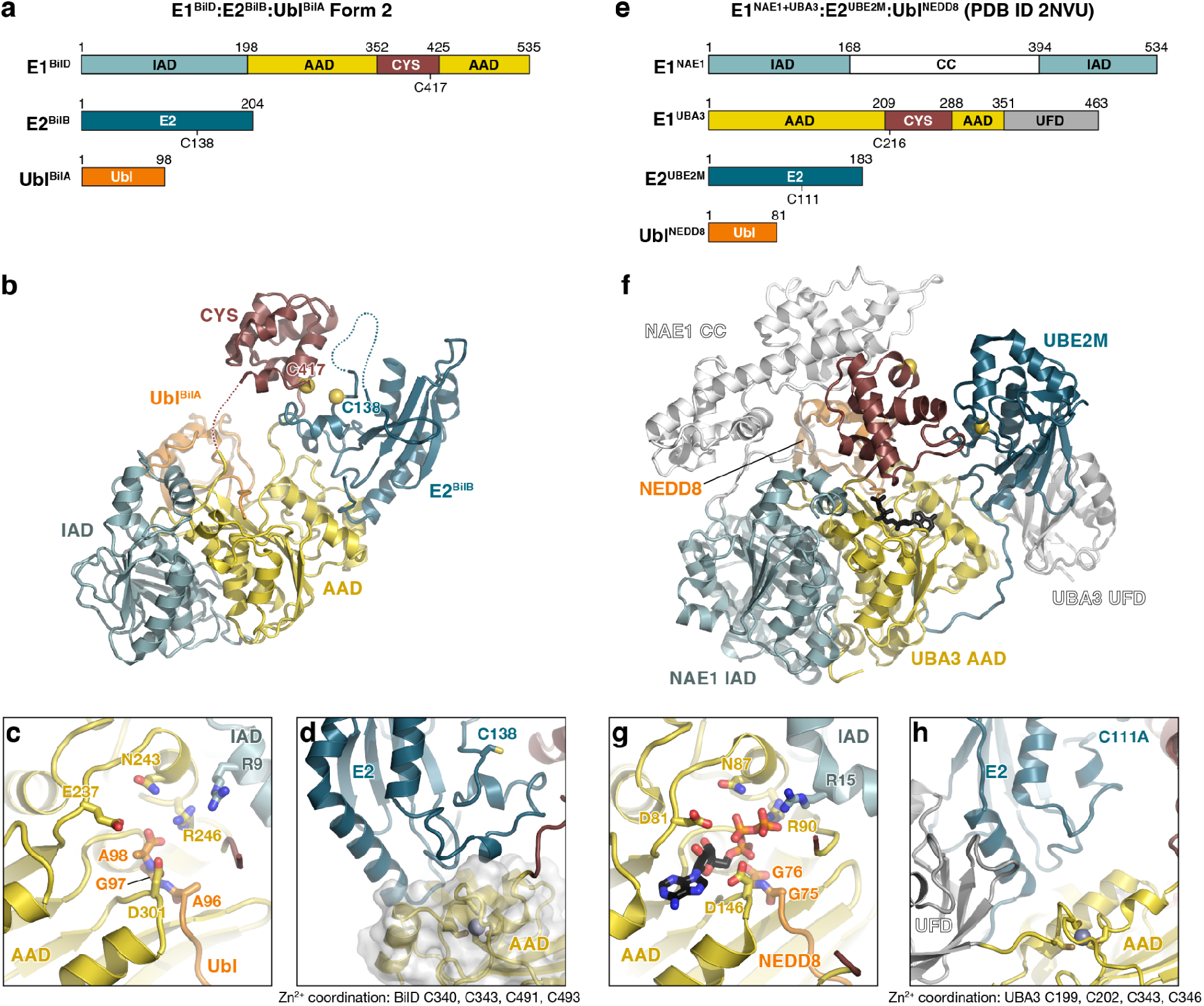
Structural parallels between bacterial and eukaryotic ubiquitination machinery. **(a)** Domain architecture of *E. aridi* E1^BilD^, E2^BilB^, and Ubl^BilA^. IAD: inactive adenylation domain; AAD: active adenylation domain; CYS: E1 catalytic cysteine-containing domain; Ubl: Ubiquitin-like domain. **(b)** Crystal structure (Form 2) of the *E. aridi* E1^BilD^:E2^BilB^:Ubl^BilA^ complex, with domains colored as in panel (a). **(c)** Closeup view of the E1^BilD^ adenylation active site with bound Ubl^BilA^ C-terminus. Conserved active site residues are shown as sticks and labeled. **(d)** Closeup view of the E1^BilD^-E2^BilB^ binding interface. **(e)** Domain architecture of *H. sapiens* E1^NAE1-UBA3^, E2^UBE2M^, and Ubl^NEDD8^ (PDB ID 2NVU)^25^. CC: coiled-coil domain; UFD: ubiquitin-fold domain. **(f)** Crystal structure of the *H. sapiens* E1^NAE1-UBA3^:E2^UBE2M^:Ubl^NEDD8^ complex (PDB ID 2NVU)^25^, with domains colored as in panel (e). **(g)** Closeup view of the E1^UBA3^ adenylation active site with bound Ubl^NEDD8^ C-terminus. Conserved active site residues are shown as sticks and labeled, and bound ATP is shown as sticks. **(h)** Closeup view of the E1^BilD^-E2^BilB^ binding interface. The UBA3 UFD is shown in gray.

**Supplementary Figure 4.**
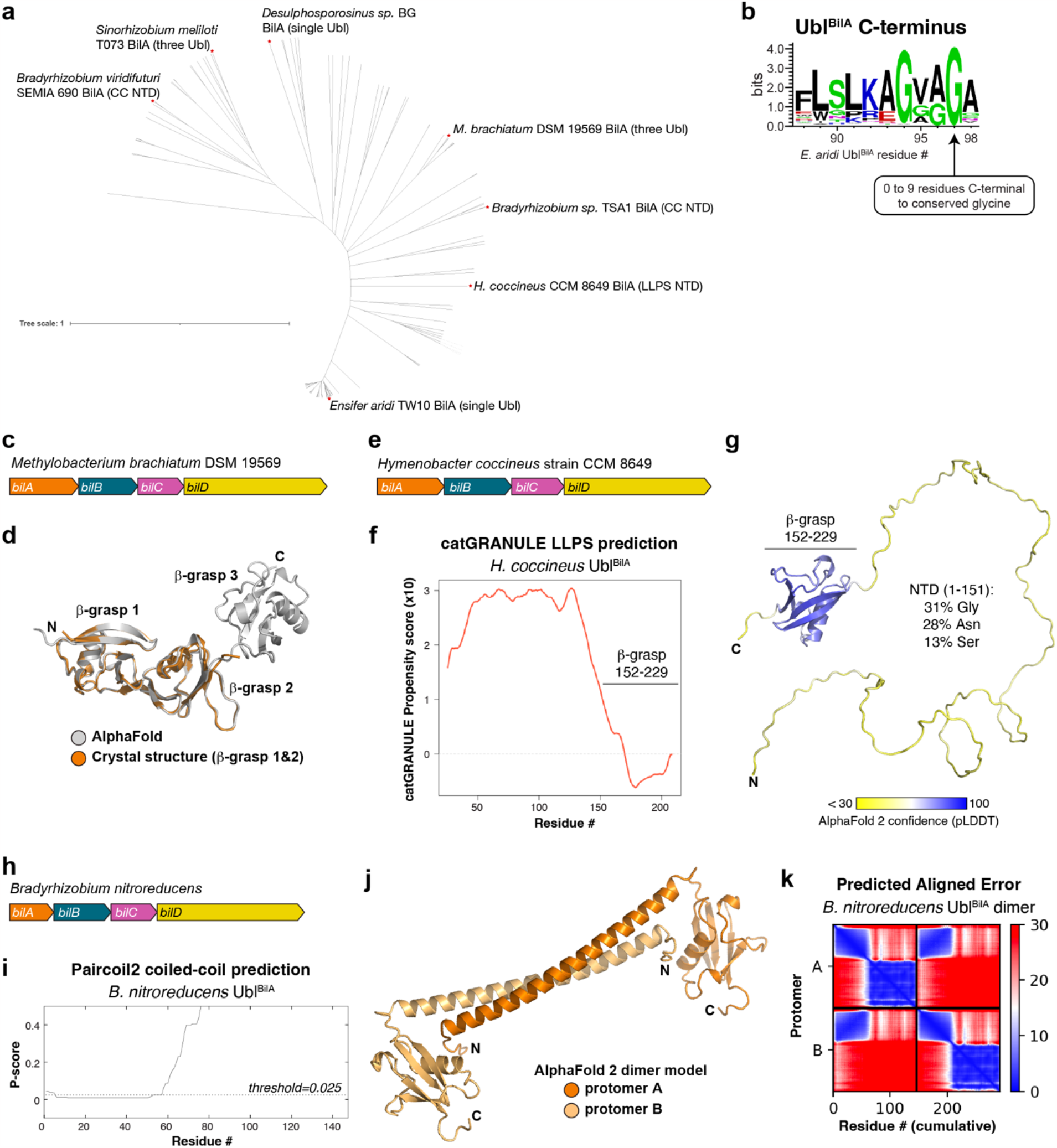
Type II BilABCD systems encode diverse Ubl proteins. **(a)** Unrooted evolutionary tree of all Ubl^BilA^ proteins in Type II Bil systems (**Supplementary Table 1**). Specific examples are labeled and their domain architectures notes. LLPS NTD: N-terminal domain predicted to undergo liquid-liquid phase separation; CC NTD: N-terminal domain predicted to form a coiled-coil. **(b)** Sequence logo from bacterial Type II Ubl^BilA^ proteins (**Supplementary Table 1**). Type II Ubl^BilA^ homologs possess up to nine residues C-terminal to the highly conserved glycine (G97 in *E. aridi* BilA). **(c)** Operon architecture of *Methylobacterium brachiatum* DSM 19569 *BilABCD* (Ubl^BilA^ IMG accession 2928979547). **(d)** Cartoon view of *M. brachiatum* Ubl^BilA^, with a full-length AlphaFold 2 model shown in gray and a 3.0 Å resolution crystal structure of β-grasp domains 1 and 2 (**Materials & Methods; Supplementary Table 1**) shown in orange. The Cα r.m.s.d. of the two structures is 0.46 Å over 134 residue pairs. **(e)** Operon architecture of *Hymenobacter coccineus* CCM 8649 *BilABCD* (Ubl^BilA^ IMG accession 2792394365). **(f)** catGRANULE^55^ profile plot showing that the N-terminus of *H. coccineus* Ubl^BilA^ is strongly predicted to undergo liquid-liquid phase separation (LLPS). **(g)** AlphaFold 2 model of *H. coccineus* Ubl^BilA^, colored by confidence (pLDDT). The disordered N-terminal domain (residues 1-151) is enriched in glycine (Gly), asparagine (Asn), and serine (Ser) residues. **(h)** Operon architecture of *Bradyrhizobium sp*. TSA1 *BilABCD* (Ubl^BilA^ NCBI accession 2881184429). **(i)** Paircoil2 ^56^ plot showing that the N-terminal domain of *B. nitroreducens* Ubl^BilA^ is predicted to form a coiled-coil. **(j)** AlphaFold 2 model of a homodimer of *B. nitroreducens* Ubl^BilA^, with one protomer colored dark orange and the second protomer colored light orange. The N-terminal domains of each protomer are arranged as an antiparallel coiled-coil dimer. **(k)** Predicted aligned error (PAE) plot for the AlphaFold 2 model shown in panel (h), showing that the N-terminal domains of each protomer are confidently predicted to interact with one another.

**Supplementary Figure 5.**
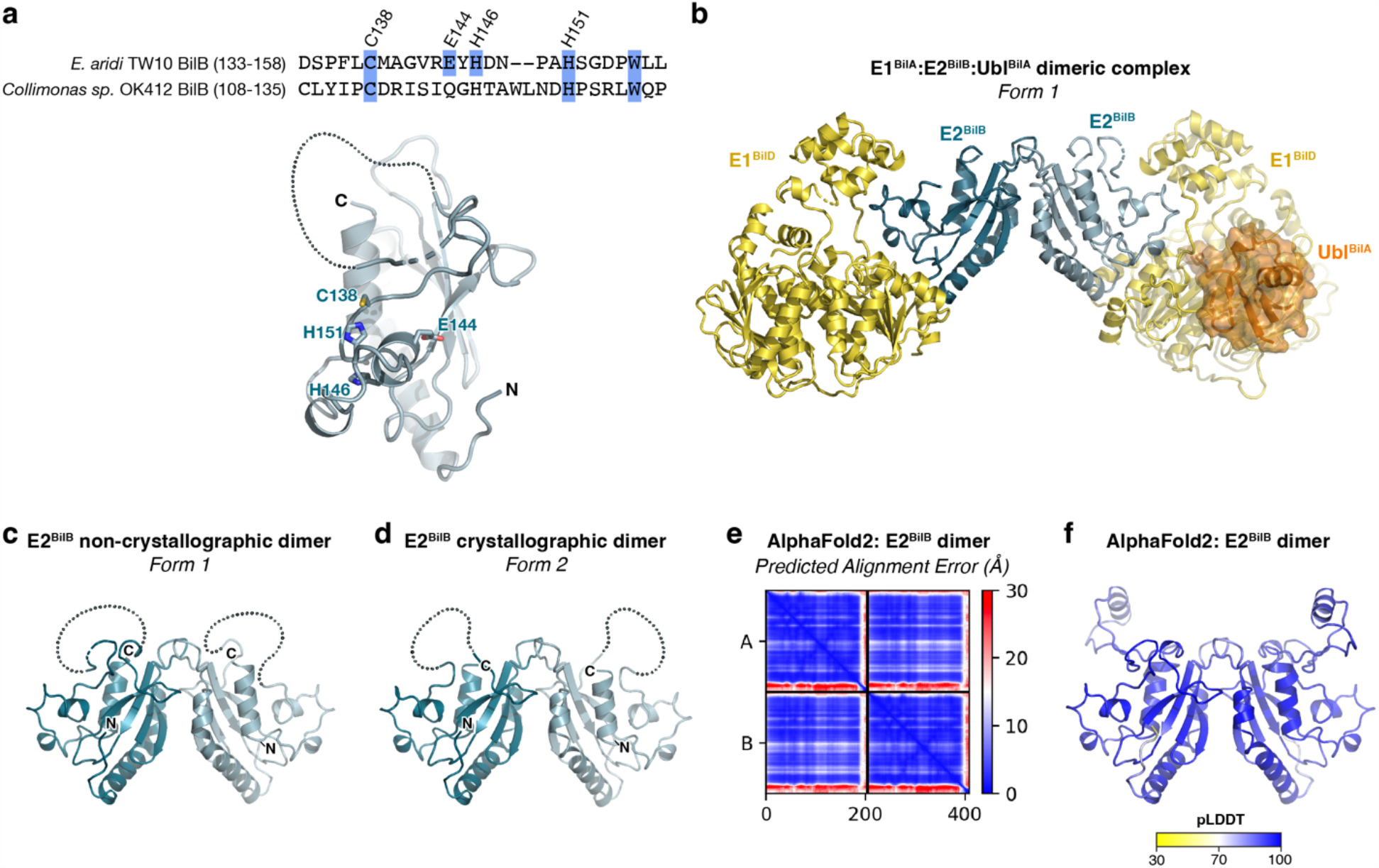
Type II Bil E2^BilB^ protein structure and homodimer formation. **(a)** *Top:* Sequence alignment of *E. aridi* E2^BilB^ and *Collimonas sp*. OK412 E2^BilB^, with the four residues of the originally identified CEHH motif in 6C/CEHH operon proteins shown in blue highlights. Of the four residues, only C138 and H151 are highly conserved across both Type I and Type II E2^BilB^ proteins. *Bottom:* Structure of *E. aridi* E2^BilB^, with the four residues of the CEHH motif shown as sticks and labeled. **(b)** Structure of a 2:2:2 E1^BilD^:E2^BilB^:Ubl^BilA^ complex, assembled through noncrystallographic symmetry in the Form 1 structure. Each subunit is colored as in Figure 1. **(c)** Closeup of the non-crystallographic E2^BilB^ dimer in the Form 1 structure. **(d)** Crystallographic E2^BilB^ dimer in the Form 2 structure. **(e)** Predicted aligned error (PAE) plot for an AlphaFold2 structure prediction of an *E. aridi* E2^BilB^ dimer. **(f)** Cartoon view of the AlphaFold2 structure prediction of an *E. aridi* E2^BilB^ dimer, colored according to confidence (pLDDT; predicted Local Distance Difference Test score). View is equivalent to panels (c) and (d).

**Supplementary Figure 6.**
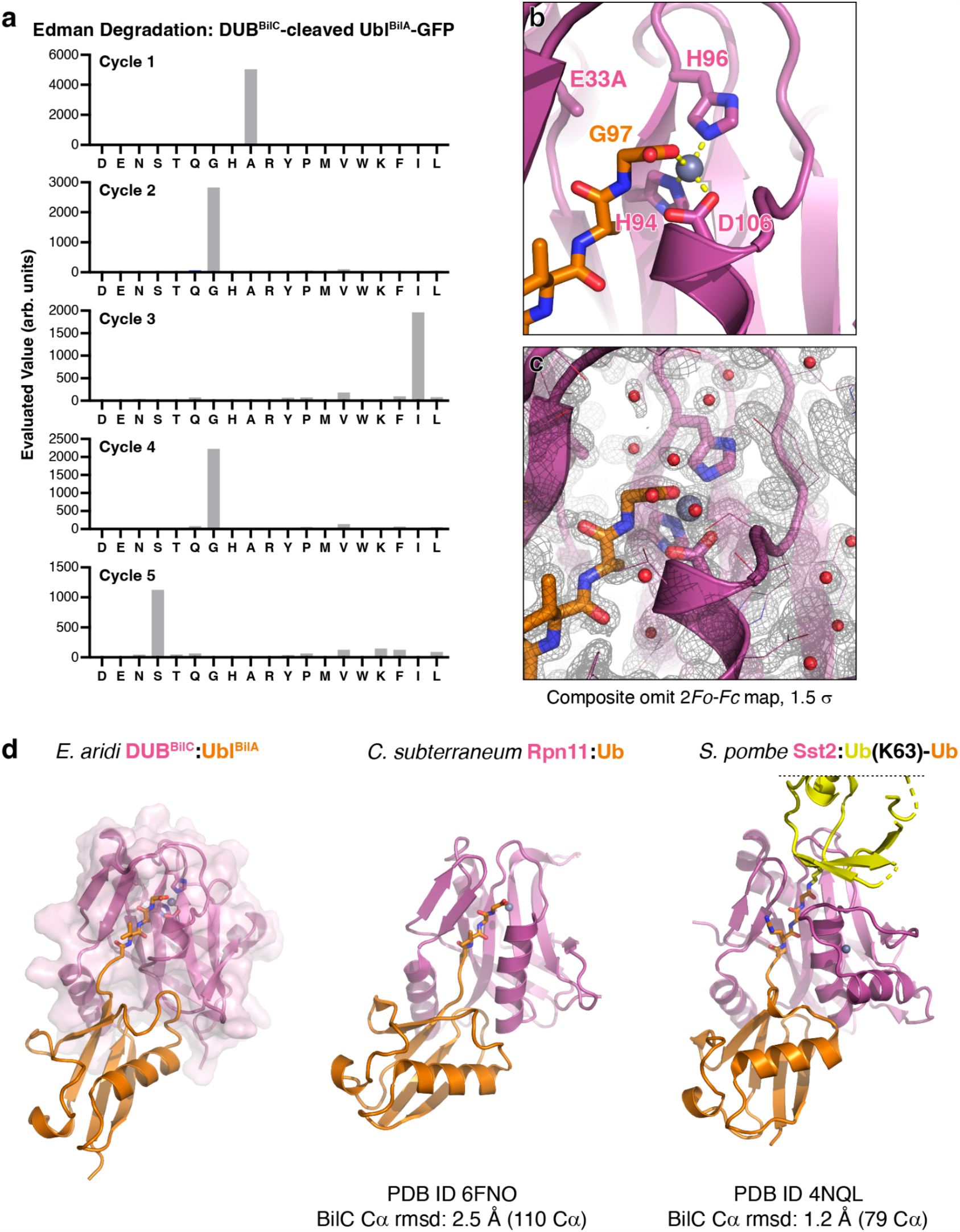
Biochemical and structural analysis of *E. aridi* DUB^BilC^. **(a)** N-terminal sequencing (Edman degradation) of DUB^BilC^-cleaved Ubl^BilA^-GFP fusion (C-terminal fragment), showing the evaluated value from each of five cycles of degradation. The inferred N-terminal sequence of the fragment is AGIGS. **(b)** Closeup view of the DUB^BilC^(E33A):Ubl^BilA^ (Form 1) active site, with proteins colored as in **Figure 3d**. Active site residues of DUB^BilC^ and glycine 97 of Ubl^BilA^ are labeled. **(c)** View equivalent to panel (b), showing 2*Fo-Fc* composite omit map density at 1.5 σ. **(d)** Comparison of *E. aridi* DUB^BilC^:Ubl^BilA^ (left) to two similar structures, *Caldiarchaeum subterraneum* Rpn11-homolog bound to ubiquitin-homolog (center)^29^ and *Schizosaccharomyces pombe* Sst2 bound to a ubiquitin K63-linked ubiquitin (right)^53^. Overall Cα r.m.s.d. values for DUB^BilC^ versus its homolog in each structure are shown.

**Supplementary Figure 7.**
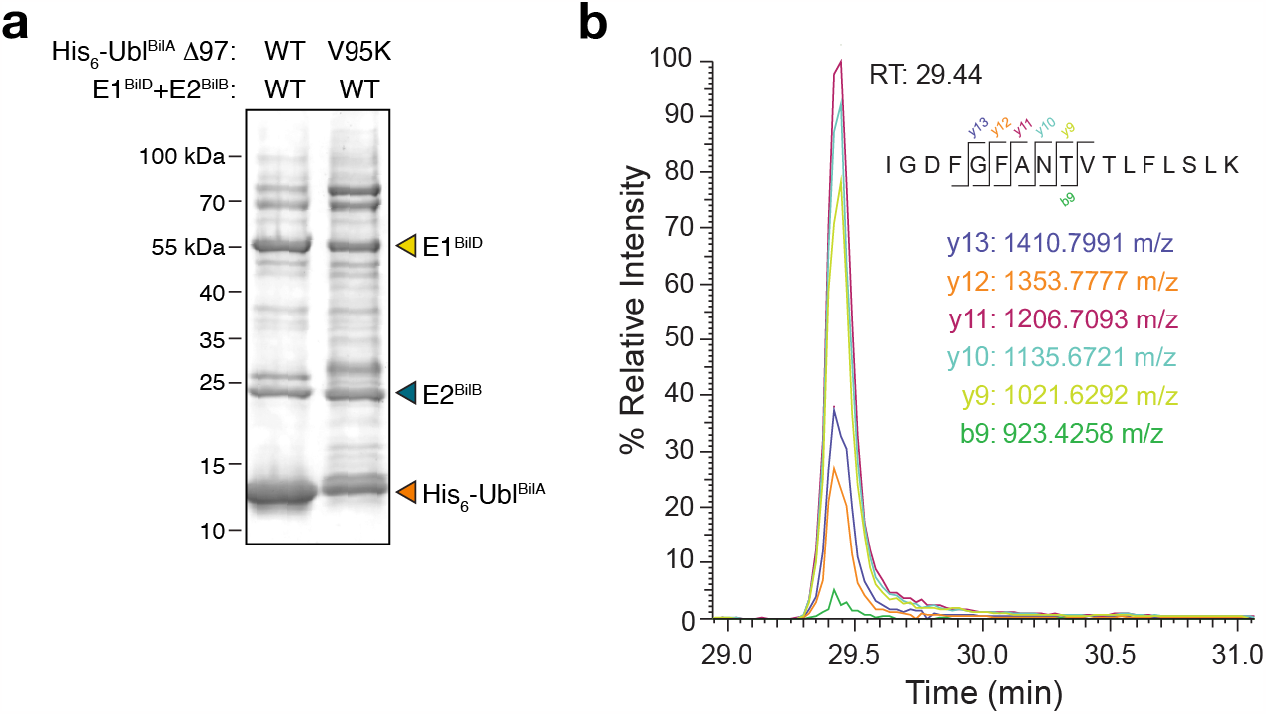
Identification of Ubl^BilA^ targets. **(a)** SDS-PAGE gel (visualized by Coomassie Blue staining) of His_6_-tagged Ubl^BilA^ coexpressed with E1^BilD^ and E2^BilB^, then purified in denaturing conditions. **(b)** Extracted Ion Chromatograms (EICs) of transition ions of the properly cleaved, unmodified Ubl^BilA^(Δ97, V95K) residues 76-92, using a 10 ppm m/z tolerance. RT: retention time.

**Supplementary Table 1. Identified Type II BilABCD operons**

*(see attached Excel spreadsheet)*

**Supplementary Table 2.**
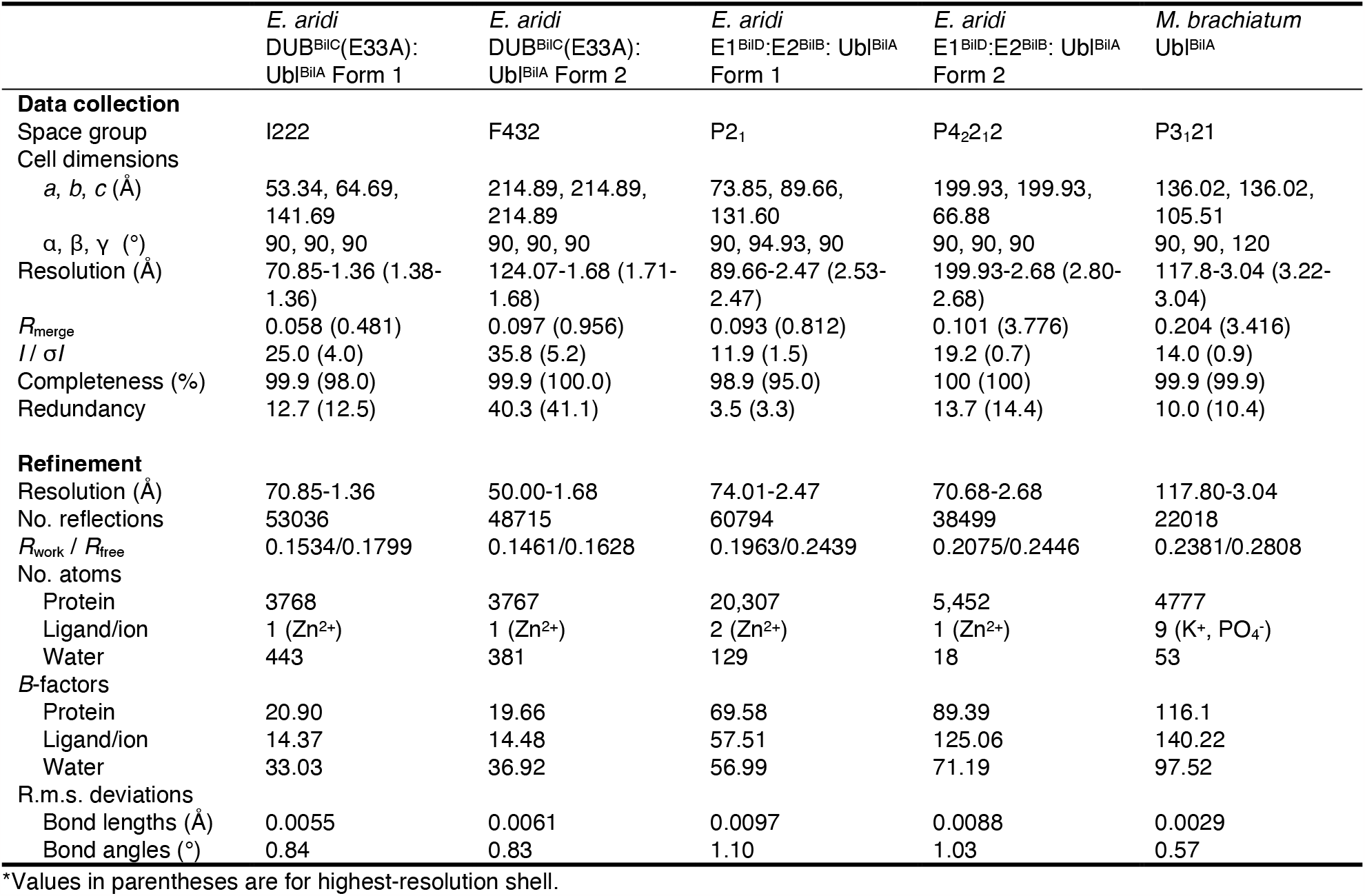
Crystallographic data collection and structure determination.

**Supplementary Table 3.**
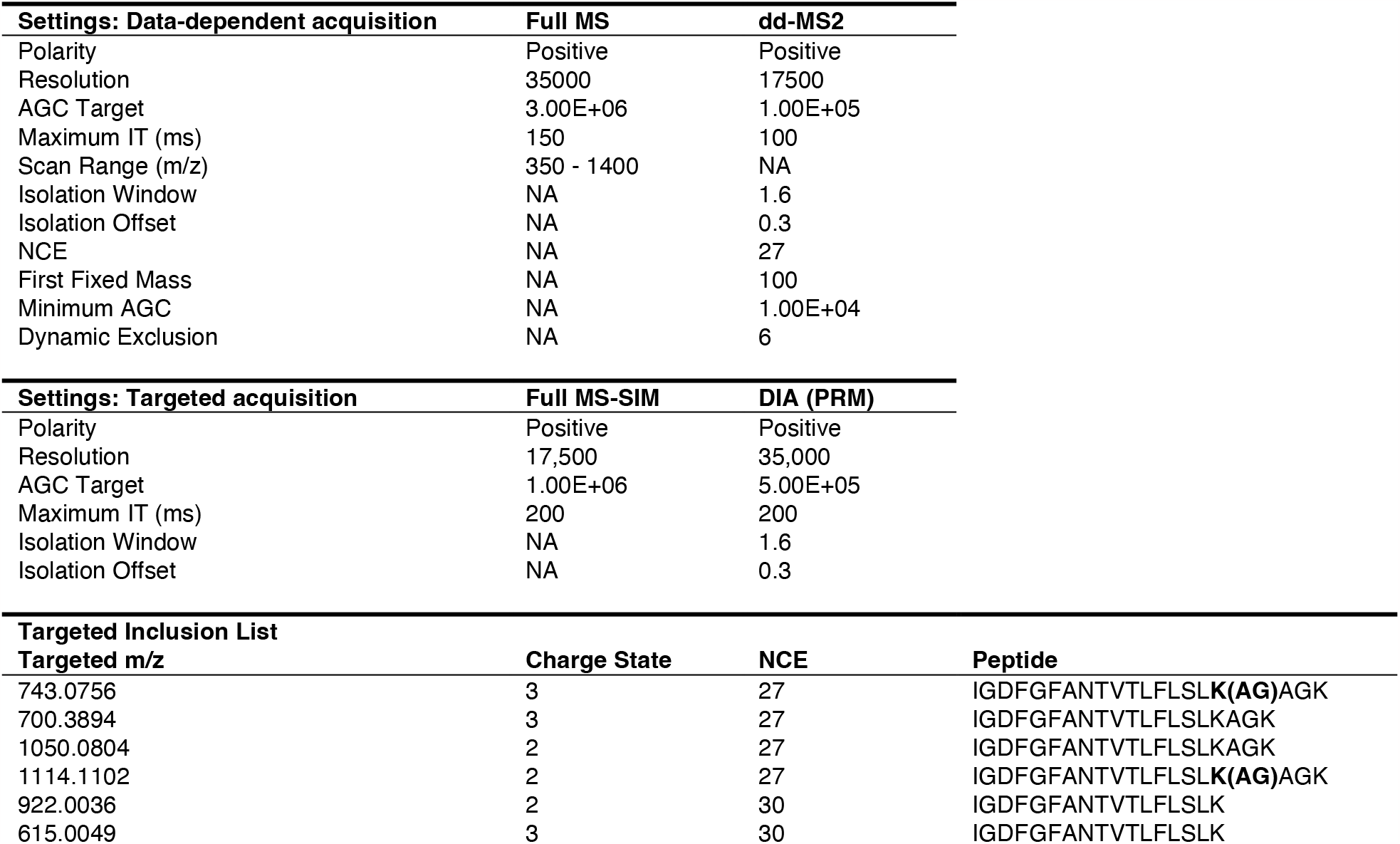
Mass Spectrometry Settings.

**Supplementary Table 4.** Peptides identified by mass spectrometry

*(see attached Excel spreadsheet)*

